# Single cell transcriptomics of vomeronasal neuroepithelium reveals a differential endoplasmic reticulum environment amongst neuronal subtypes

**DOI:** 10.1101/2024.01.08.574704

**Authors:** Devakinandan G V S, Mark Terasaki, Adish Dani

## Abstract

Specialized chemosensory signals elicit innate social behaviors in individuals of several vertebrate species, a process that is mediated via the accessory olfactory system (AOS). The AOS comprising the peripheral sensory vomeronasal organ has evolved elaborate molecular and cellular mechanisms to detect chemo signals. To gain insight into the cell types, developmental gene expression patterns and functional differences amongst neurons, we performed single cell transcriptomics of the mouse vomeronasal sensory epithelium. Our analysis reveals diverse cell types with gene expression patterns specific to each, which we made available as a searchable web resource accessed from www.scvnoexplorer.com. Pseudo-time developmental analysis indicates that neurons originating from common progenitors diverge in their gene expression during maturation with transient and persistent transcription factor expression at critical branch points. Comparative analysis across two of the major neuronal subtypes that express divergent GPCR families and the G-protein subunits Gnai2 or Gnao1, reveals significantly higher expression of endoplasmic reticulum (ER) associated genes within Gnao1 neurons. In addition, differences in ER content and prevalence of cubic membrane ER ultrastructure revealed by electron microscopy, indicate fundamental differences in ER function.

## Introduction

The Vomeronasal organ (VNO), part of the vertebrate accessory olfactory system, is a major pheromone sensing organ, that is thought to be specialized to evoke innate social behaviors in mammals. The rodent VNO neuroepithelium consists of two major neuronal subtypes, that are defined primarily based on the expression of two distinct families of G-protein coupled receptors (GPCRs) and their associated G protein alpha subunit: vomeronasal type-I GPCRs (V1R) with G⍺i2 subunit (Gnai2) neurons or vomeronasal type-2 GPCRs (V2R) with G⍺o subunit (Gnao1) neurons. V1R and V2R expressing neurons are spatially segregated within the neuroepithelium, project to distinct locations in the accessory olfactory bulb along the anterior-posterior axis. Functional evidence suggests that V1R and V2R neurons may recognize ligands that mediate different behaviors (Chamero *et al*., 2011; Oboti *et al*., 2014; Trouillet *et al*., 2019; Pallé *et al*., 2020). However, we currently lack a comprehensive mapping of receptor-ligand interactions and how these lead to distinct behaviors. Cognate ligand binding in both Gnao1 and Gnai2 neurons, is thought to signal via the phospholipase-c pathway which results in the activation of the VNO specific cation channel, Trpc2 (Liman, Corey and Dulac, 1999; Stowers *et al*., 2002; Zufall, 2005)

In the main olfactory epithelium (MOE), neurons regenerate throughout life, continuously differentiating from a stem cell population into neurons and supporting cell types (Fletcher *et al*., 2017). Neuronal differentiation is also associated with the expression of a single GPCR to the exclusion of many others (Hanchate *et al*., 2015; Tan, Li and Xie, 2015), leading to the hallmark feature termed as “one neuron one receptor” (ONOR) rule (Malnic *et al*., 1999; Serizawa *et al*., 2000, 2003). VNO sensory neurons (VSNs) also continuously regenerate (Barber and Raisman, 1978; Wilson and Raisman, 1980) from a stem cell population, and differentiating cells take on the identity of supporting cells and mature neurons of either the Gnao1 or Gnai2 sub-types (Katreddi *et al*., 2022). While Gnai2 neurons seem to largely follow the ONOR rule in expressing one V1R gene per neuron (Serizawa, Miyamichi and Sakano, 2004), Gnao1 neurons seemingly deviate from this rule by co-expressing a combination of GPCRs. Thus, each Gnao1 neuron expresses a single V2R gene from the ABD family as well as a member of the C family V2R genes (Martini *et al*., 2001; Silvotti *et al*., 2007, 2011; Ishii and Mombaerts, 2011). In addition, further molecular diversification of Gnao1 neurons occurs due to combinatorial co-expression of a family of non-classical Major Histocompatibility Complex (MHC) genes, H2-M*v*, that are co-expressed along with V2Rs (Ishii, Hirota and Mombaerts, 2003; Loconto *et al*., 2003; Ishii and Mombaerts, 2008). A few combinations of V2R and H2-M*v* expression have been identified, indicating that these co-expression patterns are non-random and could be functionally important (Martini *et al*., 2001; Ishii, Hirota and Mombaerts, 2003; Loconto *et al*., 2003; Silvotti *et al*., 2007, 2011; Ishii and Mombaerts, 2008, 2011). However, identifying the precise combinatorial expressions of V2R-ABD, V2R-C and H2-M*v* has been elusive, partly due to the challenges in generating RNA or antibody reagents to identify closely homologous genes, combined with the challenges with multiplexing more than three targets.

Starting with a common progenitor population, how do cells of the VNO sensory epithelium diversify in their gene expression patterns that define the sensory neuron types? Recent studies using single cell transcriptomic analysis have implicated Notch signaling along with downstream transcription factors, Tfap2e, Bcl11b to play a role in specifying the fates of developing VSNs (Enomoto *et al*., 2011; Katreddi *et al*., 2022; Lin *et al*., 2022). As the progenitor cells differentiate to mature neurons, transcriptional reprogramming downstream to Notch signaling leads to distinct neuronal populations: Gnao1 and Gnai2 neurons. While these studies identified transcription factors, it is not clear whether the two major differentiated VSN subtypes expressing Gnao1-V2Rs or Gnai2-V1Rs fundamentally differ in their cellular properties.

Here we applied a multilevel transcriptomics approach which bundles single cell RNA sequencing (scRNA seq) validated by low-level spatial transcriptomics to characterize gene expression of the mouse VNO sensory epithelium. Our results identified genes specifically expressed in sensory, non-sensory cell types and the divergence of gene expression between Gnao1 - Gnai2 neurons at different maturation stages. Analysis of mature neurons provided a comprehensive picture of specific co-expression patterns of VR and H2-M*v* genes. Surprisingly, mature Gnao1 neurons revealed a significant enrichment of endoplasmic reticulum related gene transcripts and proteins, which is indicative of a fundamental divergence in cellular function between these cell types. Electron microscopy further revealed a hypertrophic ER ultrastructure consisting of gyroid cubic membrane morphology, more prevalent in Gnao1 neurons.

## Results

### Single cell transcriptome of mouse vomeronasal neuroepithelium

We standardized VNO neuroepithelium dissociation to obtain a single cell suspension along with optimizing cell viability. We performed single cell droplet-based capture coupled with scRNA seq, using the 10X genomics platform on four biological replicates consisting of male and female animals. Initial quality control steps were performed (see methods) to remove cells with high or low total RNA content and red blood cell contamination, resulting in the retention of 9185 cells from all samples for further analysis. Since our comparison of gene expression obtained from male versus female mice, did not reveal appreciable differences other than known sexually dimorphic genes that include Eif2s3y, Ddx3y, Uty and Kdm5d (**Figure 1-figure supplement 1, supplementary table 1)**, we pooled cells from both sexes for further analysis.

To group cells based on their gene expression profiles, we performed dimensionality reduction and unsupervised clustering, which resulted in 18 distinct clusters (**Figure 1A**), with gene expression markers specific to each cluster (**Figure 1B**). Major markers for each cluster were identified by differential expression analysis and the cell type identity was assigned to each cluster based on known markers for each cell type. Amongst 9185 cells, we observed that 63.7 percent (5852 cells) were neuronal cell types at different developmental stages which includes progenitor cells (Cluster 5; defined by expression of Ascl1, Neurod1, Neurog1), immature neurons (Cluster12; expression of Gap43), Gnao1 neurons (Cluster 14; based on expression of Gnao1, V2Rs), and Gnai2 neurons (Cluster 7, 16; characterized by Gnai2, V1Rs expression) (**Figure 1C**). The remaining 36.3 percent cells were classified to be of non-neuronal types which include sustentacular cells (Cluster 15; defined by Cyp2a5, Ppic), cells from non-sensory epithelium (Cluster 11; defined by Ly6d, Krt5), cells from glandular tissue (Cluster 13; characterized by Obp2a, Lcn3), immune cells which include macrophages (Cluster 8, 6, 10; defined by C1qa, C1qb, H2-Aa), T cells (Cluster 9; defined by Cd3d, Cd3g), neutrophils (Cluster 2; defined by s100a8, s100a9), olfactory ensheathing cells (Cluster 3; defined by Gfap, Plp1, Sox10), endothelial cells (cluster 1 and 17; defined by Gng11, Egfl7, Eng), fibroblasts (Cluster 4; defined by Igfbp4, Dcn, Lum) and solitary chemosensory cells (Cluster 18; defined by Trpm5, Gnat3). A complete list of top 20 differentially expressed genes associated with each cluster and proportion of cells are provided in **supplementary table 2 and 3**.

**Figure 1.**
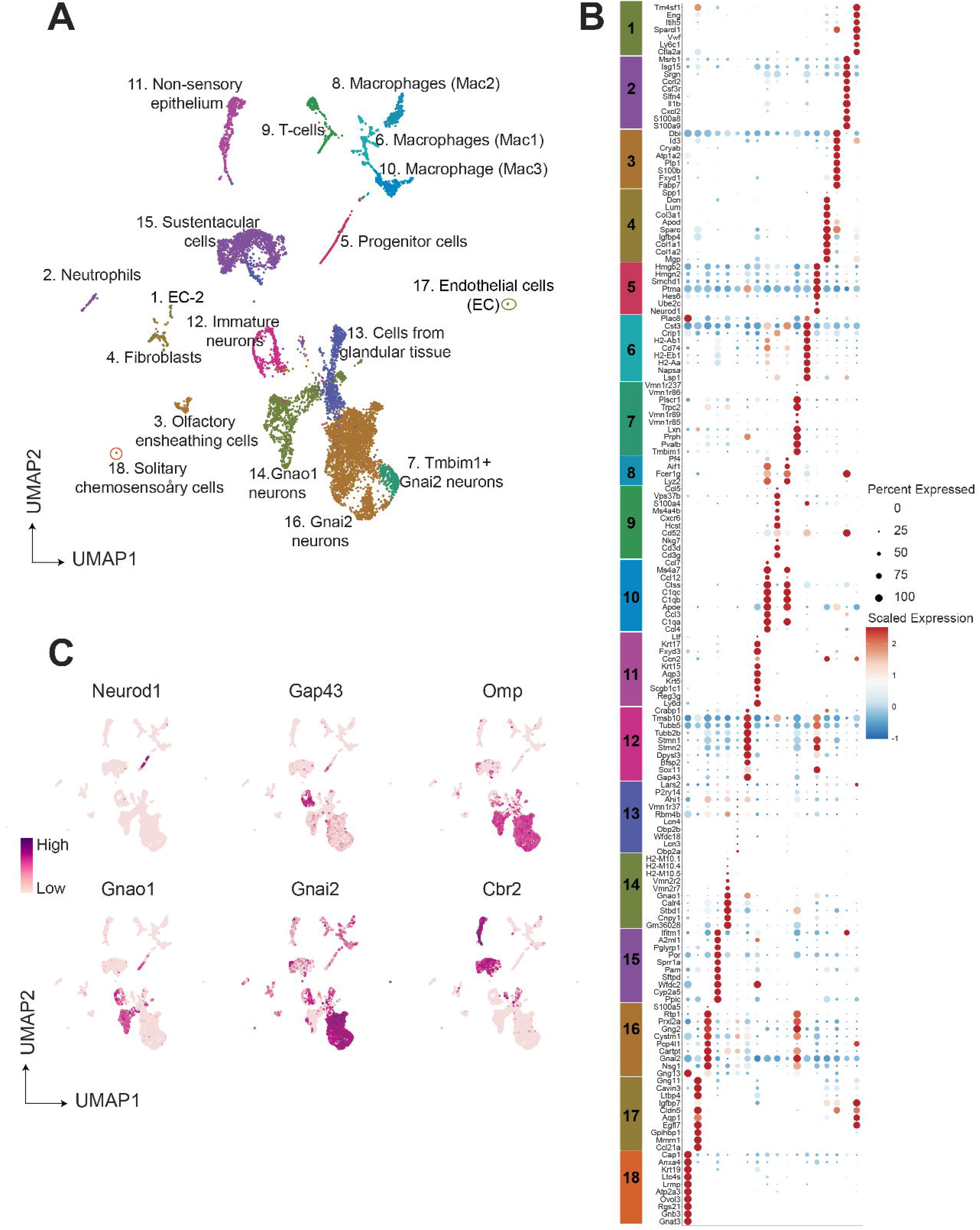
Single cell RNA sequencing of the mouse vomeronasal neuroepithelium **A)** Uniform Manifold Approximation and Projection (UMAP) of 9185 cells from vomeronasal neuroepithelium with 18 identified clusters. Each point represents a cell that is color coded according to the cell type. Clusters were assigned a cell type identity based on previously known markers. Percentage of total cells corresponding to each cluster is in **Supplementary table 3. B)** Dot plot showing the expression of top 10 gene markers for each cluster identified by differential expression analysis. Gene expresion markers shared across multiple clusters are listed only once. The dot radius and dot color indicate the percentage of cells expressing the gene and scaled expression value in that cluster. respectively. Top 20 gene markers for each cluster with log2 fold change are listed in **Supplementary table 2. C)** Feature plot showing the expression level of major known gene markers of neuronal and sustentacular cell types.

We performed RNA *in situ* hybridization (RNA-ISH) on VNO sections to confirm spatial classification of various non-sensory cell type clusters based on some of the genes identified in our study (**Figure 2A**). Thus, we confirmed Ppic to be expressed in sustentacular cells and Ly6d in cells of non-sensory epithelium bordering the VNO lumen. Since, Cbr2 - a known marker of sustentacular cells is also expressed in non-sensory epithelium (**Figure 1C**), Ppic and Ly6d are better markers to distinguish them. Obp2a was confirmed to be expressed in cells from glandular tissue on the non-sensory side. Plp1 and Mgp, markers of olfactory ensheathing glial cells and endothelial cells/fibroblasts, respectively, were observed to be expressed in cells located at the periphery of the neuro-epithelium. Macrophage cell markers C1qb and H2-Aa were observed to be specifically expressed in scattered cells throughout the neuroepithelium, with H2-Aa co-expression also seen in sustentacular cells **(Figure 2A)**, confirming the presence of these cell types within the neuroepithelium.

**Figure 2.**
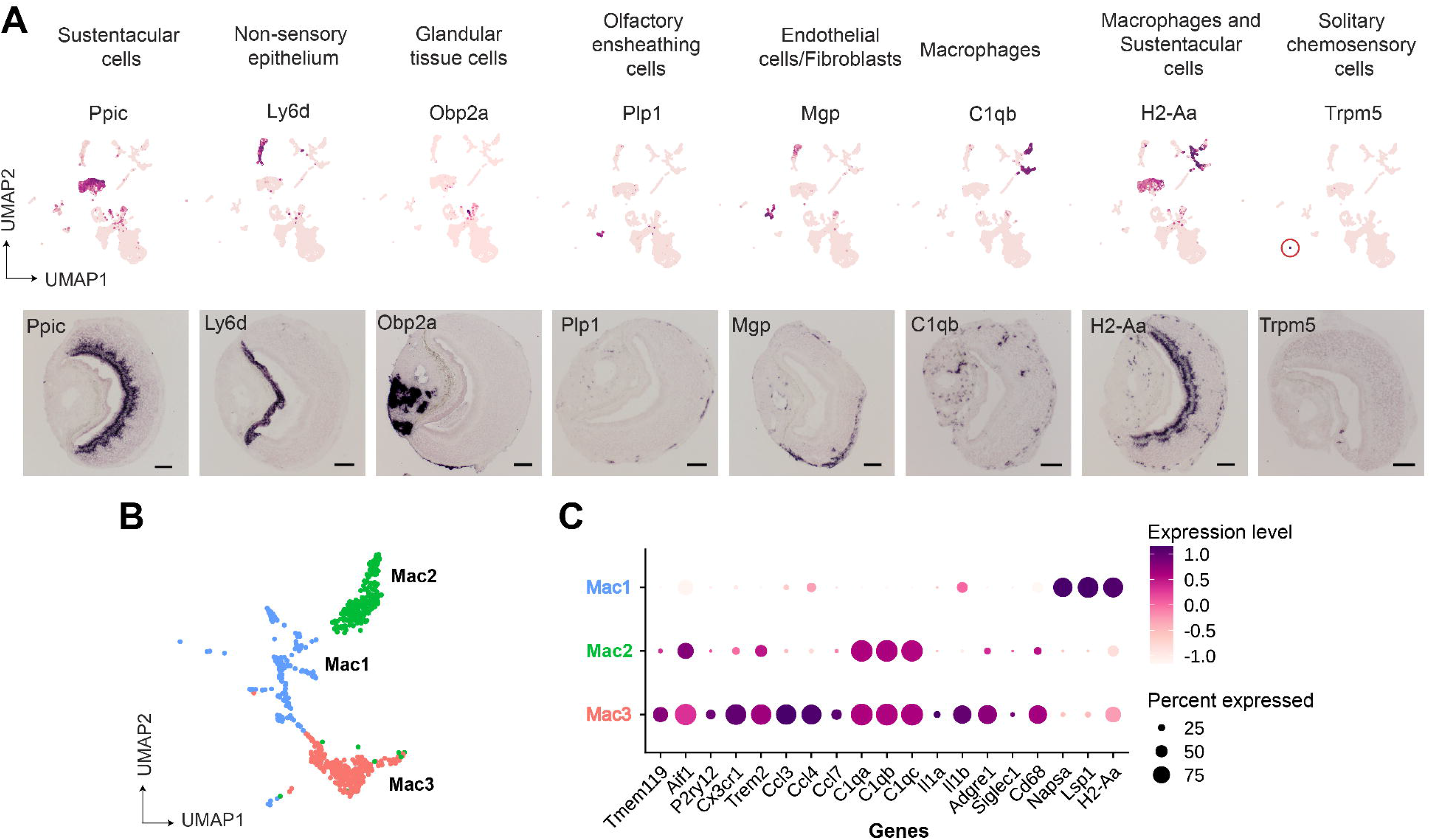
Gene markers of non-sensory cells in the vomeronasal organ identified from single cell RNA sequencing. A) UMAP feature plot (top) of a single representative gene marker for non-sensory cell clusters and their spatial gene expression pattern by RNA *in situ* hybridization on VNO coronal sections (bottom, scale bar: 100µm). **B, C)** Three custers of macrophage-like cell types represented by UMAP projection **(B)** and a dot plot **(C)** showing differ­ entially expressed genes between these clusters. The dot radius indicates percentage of cells expressing a gene, whose scaled expression level is represented by intensi­ ty of color as per the scale. Additional markers of each non-sensory cell type are in Supplementary table 1 and differential expression between macrophage clusters is in **Supplementary table 4.**

Since our study identified multiple immune cell clusters, we examined these in some detail. Three clusters (**Figure 1A**, clusters 6, 8, 10 termed as Mac1, Mac2, Mac3 respectively) comprised of macrophage like cells based on expression of C1qa, C1qb, C1qc and MHC class II gene - H2-Aa **(Figure 2B, 2C)**. Further analysis indicates that cluster 10 (Mac3) expresses classical microglial markers such as Tmem119, Aif1 (Iba1), P2ry12, Cx3cr1, Trem2, along with chemokine genes - Ccl3, Ccl4 and Ccl7 and Interleukins – Il1a, Il1b (**Figure 2C)**, indicating an activated macrophage subset. Cluster 8 (Mac2) is similar to Mac3 in regard to expression of classical tissue resident macrophage marker - Adgre1 (F4/80), C1qa, C1qb, C1qc but lacks expression of chemokine and interleukin genes mentioned earlier. In comparison to Mac3 and Mac2, the Mac 1 cluster is distinguished by the lack of C1qa, C1qb, Cd68, Adgre1(F4/80), microglial markers but possesses high levels of Napsa, Lsp1, H2-Aa **(Figure 2C, Supplementary table 2, 4)**. In summary, the identification of discrete macrophage cell types from our data, along with experimental validation of their presence using some of the expression markers, points to the possibility of tissue resident or tissue-infiltrating macrophages within the VNO neuroepithelium. These VNO macrophage types could have functional implications towards influencing processes such as neuronal differentiation or responses to tissue repair and injury.

On the other hand, we did not observe evidence via RNA-ISH for T-cell markers (Cd3d, Cd3g), raising the possibility that these immune cell types may have co-purified from blood. Lastly, we were able to identify via RNA-ISH, expression of Trpm5 **(Figure 2A)** and Gnat3 genes from our dataset, that mark solitary chemosensory cells expressing taste receptors and related markers (Ogura *et al*., 2010) (**Supplementary table 2**), indicating that rare cell-populations were captured, and their markers identified in our study.

### Neuronal cell types and transient gene expression in developing neurons

Our next step was to perform in depth analysis of vomeronasal sensory neurons. In order to obtain a better resolution on neuronal sub-types, We pooled cells from clusters representing neuronal types identified in Figure-1 (clusters 5, 7, 12, 14, 16), created a new Seurat object and performed re-clustering. This resulted in 13 clusters (labelled n1- n13), which potentially represent distinct neuronal sub-populations within the broadly defined Gnao1 and Gnai2 neurons (**Figure 3-figure supplement 1A**). Since neurogenesis is a continuous process in VNO neurons (Barber and Raisman, 1978; Wilson and Raisman, 1980), these clusters also represent a snapshot of cells at different developmental states, identified by known markers: Globose basal cells (Cluster n5; Ascl1), progenitor cells (Cluster n5; defined by expression of Neurod1, Neurog1), immature neurons (Cluster n6, n7; defined by Gap43) expressing Gnao1 or Gnai2 (**Figure 3-figure supplement 1B**). Among Gap43 negative mature neurons, Gnao1 neurons are clusters n1-n4, (**Figure 3-figure supplement 1A, B**, corresponding to cluster 14 in **Figure 1A**), where sub-cluster identity is clearly defined by their co-expression of H2-Mv and Vmn2r GPCRs (discussed in detail in the co-expression section below). In contrast, 6 clusters (n8-n13) correspond to mature Gnai2 neurons (clusters 7, 16 from **Figure 1A**), of which n8-n11 do not have any definite gene markers and differ based on differences in the expression level of few genes (**Figure 3-figure supplement 1C, Supplementary table 5**). Neurons of sub-cluster n12 (cluster 7 from **Figure 1A**) exclusively express genes Tmbim1, Lgi2, Cleg2g and n13 are distinguished by the expression of previously identified activity dependent genes - Rasgrp4, Pcdh10 and S100a5 (Fischl *et al*., 2014) (**Figure 3-figure supplement 1C, Supplementary table 5**. To our knowledge, unlike mature Gnao1 neurons, mature Gnai2 subclusters n8-n11 and n13, do not have an obvious principal parameter for their subclassification.

**Figure 3.**
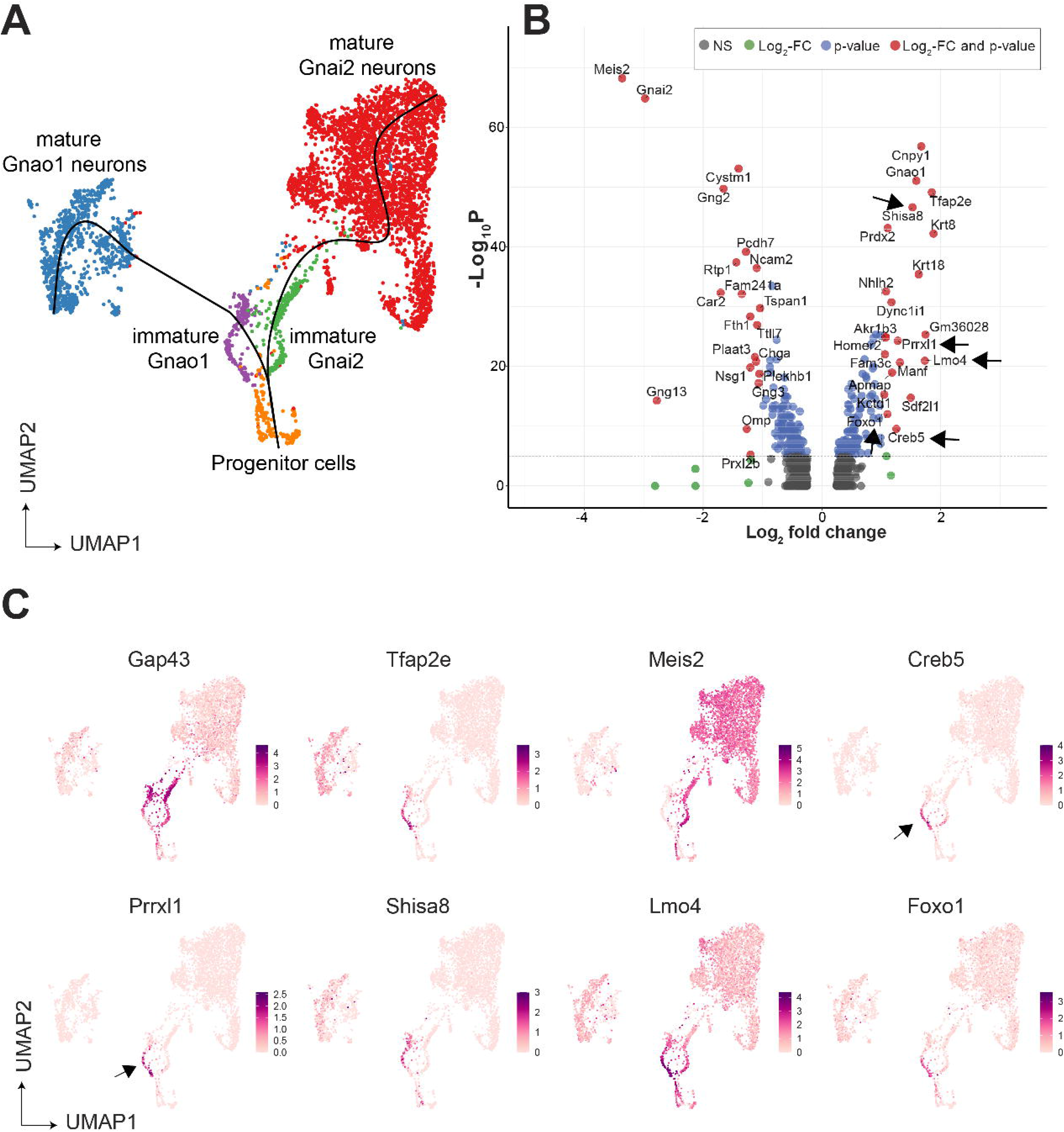
Gene expression dynamics during development of sensory neurons **A)** UMAP of neuronal cells with clusters (n1-n13 from Figure 3, Figure 3-figure supplement-1) represented in different colors. Sub-clusters within mature Gnai2, Gnao1 neurons are merged_ The line on the UMAP plot shows pseudotime developmental trajectory of neurons. **BJ** Volcano pot of differential gene expression between immature Gnai2 (cluster n6; Gap43+,Gnai2+) vs immature Gnao1 (cluster n7; Gap43+,GnaO1+) neurons_ Genes that satisfy 1109_2_ fold change normalized expression! > 1 (green) and -109_10_ p value > 6 (blue) are considered significant and coloured in red_ Non-significant (NS) genes are colored in grey_ Arrows point to transcription regulators enriched in Gnao1+ immature neurons. Complete list of differentially expression genes is in Supplementary table 6. CJ Feature plots showing the normalized expression levels of previously known markers for immature neurons (Gap43), Gnao1 neurons (Tfap2e), Gnai2 neurons (Meis2) and transcription factor or related genes: Creb5, Prrxl1, Shisa8, Lmo4, Foxo1_Arrows highlight the limited expression of Creb5 and Prrxl1 to immature neurons, but absent from mature indicating transient expression during development of Gnao1 neurons. Subclusters within Gnao1, Gnai2 neurons and effect of VRs on subclustering are in Figure 3**-figure supplement 1, 2** respectively.

Expression of cognate V1R or V2R GPCRs is one of the hallmarks of VSN differentiation. We asked the question, to what extent do VRs influence neuronal sub clustering. Exclusion of VRs from clustering parameters did not affect divergence of mature neurons into Gnai1 / Gnao1 and only selectively affected some of the subclusters (**Figure 3-figure supplement 2)**, thus indicating that neuronal subtype identity was governed by gene expression differences beyond vomeronasal GPCRs.

Since immature neurons separated into two clusters expressing Gnao1 or Gnai2, these imply a possible branch point in the developmental trajectory towards mature neurons. To confirm this, we performed pseudo time analysis on neurons using Slingshot, and progenitor neurons (cluster n5) as a start/anchor point. For the purpose of developmental analysis, we merged subclusters within mature Gnao1 and Gnai2 neurons. The trajectory of development confirms a split of immature Gap43+ neurons into Gnao1 or Gnai2 cell clusters (**Figure 3A**). We performed differential expression analysis between immature neurons (cluster n6: Gap43+, Gnao1+ vs cluster n7: Gap43+, Gnai2+) (**Figure 3B, Supplementary Table 6**). To identify potential transcription factors in this developmental branch point, we manually curated genes that have molecular features associated with DNA binding domains or transcription regulation based on literature (**Figure 3C**). To our surprise, we found multiple genes that are known transcription regulators but are not previously reported to be enriched in Gnao1 neurons. These include Creb5, Prrxl1, Shisa8, Lmo4 and Foxo1. Creb5, Prrxl1 are transcription factors expressed only in immature Gnao1 neurons whereas Shisa8, Lmo4 and Foxo1 expression is enriched in immature Gnao1 neurons compared to mature ones (**Figure 3C**). Prrxl1 was reported to auto-repress its expression which explains how its expression is limited to immature Gnao1 neurons (Monteiro *et al*., 2014).These data indicate that the developmental transition of VNO neurons to Gnao1 lineage may involve the transient expression of transcription factors within immature neurons after dichotomy is established.

### Co-expression of Vomeronasal receptors

One of the features that differentiates vomeronasal sensory neurons from main olfactory sensory neurons is their systematic deviation from ‘one neuron one receptor rule’, especially in mature Gnao1 neurons. V2R receptors have been classified into families- A, B, D, E and a distinct family-C that is phylogenetically closer to calcium sensing receptor (Young and Trask, 2007). Several studies have established a broad pattern of co-expression amongst these V2R family members, such as members of family ABDE co-express with family-C GPCRs and components of family-C (Vmn2r1 -also termed - C1, Vmn2r2-Vmn2r7 -termed C2) in turn co-express with each other in specific combinations (Martini *et al*., 2001; Silvotti *et al*., 2007). Furthermore, H2-M*v* genes within Gnao1 neurons also exhibit non-random co-expression patterns with the V2R GPCRs. (Ishii, Hirota and Mombaerts, 2003; Loconto *et al*., 2003; Silvotti *et al*., 2007, 2011; Ishii and Mombaerts, 2008, 2011). To further investigate these co-expression patterns and subpopulations within Gnao1 neurons, we took a closer look at clusters n1-n4 comprised of mature Gnao1 neurons (**Figure 3-figure supplement 1A)**. These four clusters **(Figure 4A)** are found to be organized majorly based on the expression of family-C V2Rs and H2-M*v* genes, especially Vmn2r1 or Vmn2r2 (**Figure 4B-E**). Of the 4 clusters, cluster n1 and cluster n4 express Vmn2r1 (**Figure 4B, 4E**) whereas cluster n3 neurons are distinguished by the co-expression of Vmn2r2 along with other family-C members (Vmn2r3, Vmn2r4, Vmn2r5, Vmn2r6. Vmn2r7) (**Figure 4C, 4E**). Cluster n2 is a mixture of either Vmn2r1 or Vmn2r2 expressing neurons (**Figure 4E**), with Vmn2r2 being similarly co-expressed along with other family-C members in those cells. We observed that among family-ABDE members, A1-A6 V2Rs are highly expressed in cluster n2, n3 which co-express Vmn2r2 whereas A8-A10, B, D, E family V2Rs are mainly expressed along with Vmn2r1 in cluster n1 neurons **(Figure 4E)**. These observations of family biased expression of ABDE with Vmn2r1 or Vmn2r2 agrees with earlier studies (Silvotti *et al*., 2007, 2011) that have used antibody or RNA probe based approaches.

**Figure 4.**
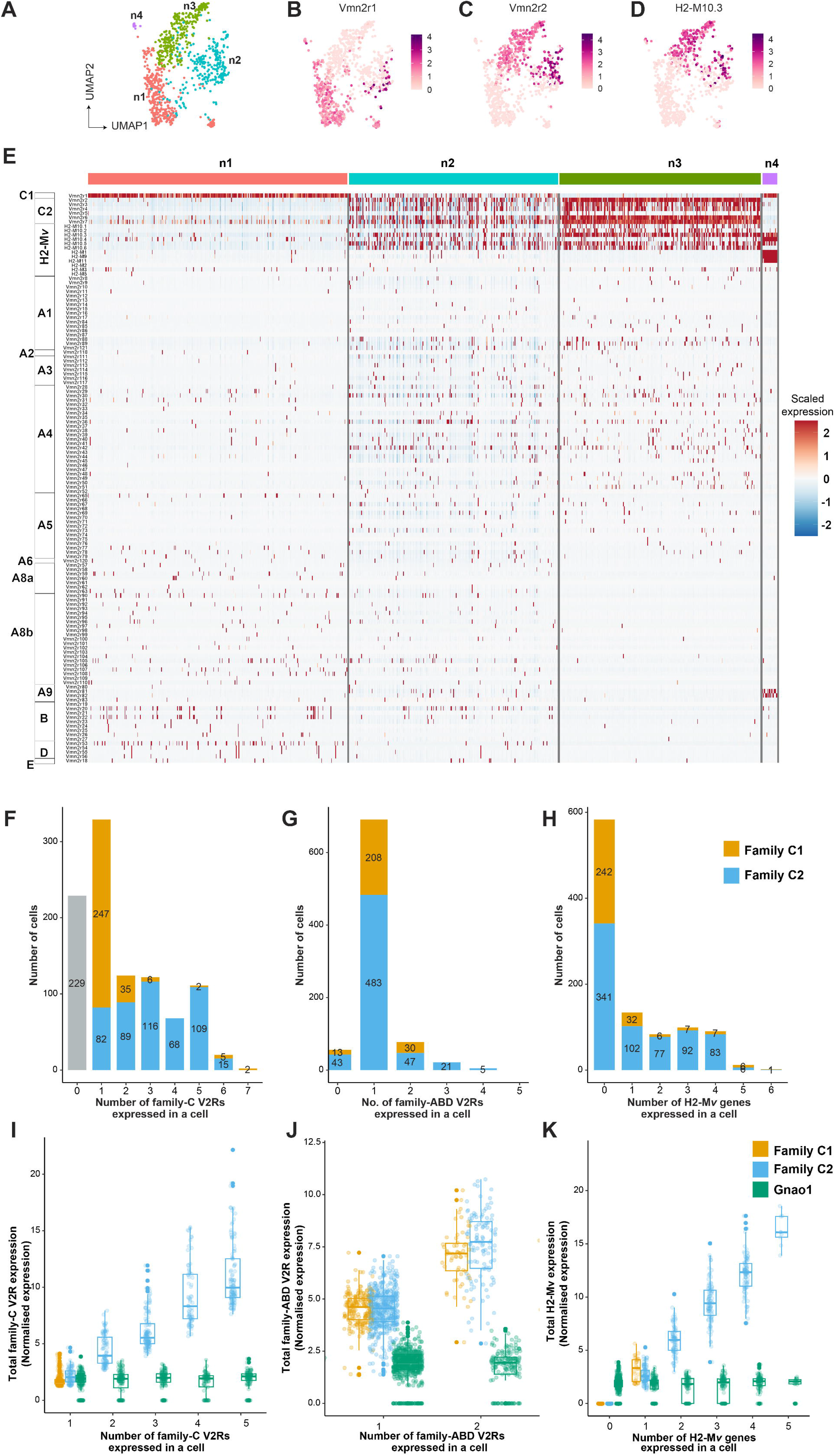
Subpopulation of Gnao1 neurons defined by V2R and H2-Mv expression. A) UMAP representation of Gnao1 neurons from Figure 3. Each dot represents a cell and four Gnao1 neuron clusters (n1-n4) are marked in different colours. **B-D)** Feature plot showing exclusive expression of Vmn2r1 **(B),** Vmn2r2 **(CJ** and the most abundant H2-Mv gene, H2-M10.3 **(DJ** in Gnao1 neurons. **E)** Heat map showing the expression of V2R and H2-Mv genes in Gnao1 neurons. Cluster numbers are marked on the top with color coding as in **(A)** and gene families are labelled on the left. Each column represents a cell and the scaled gene expression in each row is colour coded as per the scale with red and blue indicating high and low expression respectively. Vmn2r1 is expressed in almost all cells of cluster-1 and cluster-4; Vmn2r2 is expressed in all cells of cluster 3; Cluster2 has mutually exclusive expression of Vmn2r1 and Vmn2r2. **F-H)** Bar plot showing number of cells expressing: 0-7 family-C V2Rs per cell **(F),** 0-5 family-ABD V2Rs per cell **(G),** 0-6 H2-Mv genes per cell **(H)** with composition of cells associated with family C1 (orange) or C2 (blue) V2R color coded on the bar. I) Box plots comparing the sum of normalised expression levels of family-C V2Rs and Gnao1 (Green) in a cell that expresses 1 to 5 fami­ ly-C V2Rs. **J)** Box plot comparing the level of total ABD-V2R expression from cells with 1 and 2 ABD-VRs along with Gnao1 expression level (green). **K)** Box plot comparing the level of total H2-Mv expression in cell that express 1-5 H2-Mv genes along with Gnao1 expression level (Green). Multiple combinations of family-C, family ABD V2Rs and H2-Mvs identified to be co-expressed in a single cell and their cell frequency are listed in Supplementary table 7.

Furthermore, we observed that expression of H2-M*v* genes (H2-M10.1, M10.2, M10.3, M10.4, M10.5, M10.6) is largely confined to clusters n2 and n3, where they are co-expressed selectively in Vmn2r2 positive neurons and mostly absent from Vmn2r1 expressing cluster n1, n2 neurons (**Figure 4D, 4E,** **Figure 4–figure supplement 2A**). These observations align with prior reports from antibody or RNA probe-based experiments, that demonstrated a sub-division amongst Gnao1 neurons based on their expression of family-C Vmn2rs and co-expression of H2-M*v* genes with the Vmn2r2 subset (Silvotti *et al*., 2007).

We then asked whether at the single neuron level, we could identify trends in co-expression patterns between ABD-family V2Rs and H2-M*v* genes, or between members of ABD family themselves. We performed co-expression analysis to identify specific VR and H2-M*v* combinations by setting an expression threshold from normalized counts. The distribution of normalized expression for VRs and H2-M*v*s across cells was bimodal, with the first peak near zero and a second peak likely corresponding to true expression (**Figure 4-figure supplement 1D-F**). We set starting of second peak as a threshold to call the expression of VRs or H2-M*v*s in a single cell. Thus, a VR or H2-M*v* was considered as co-expressed in a cell only if its normalized expression value surpasses the set threshold based on the distribution (**Figure 4-figure supplement 1E, 1F**). The threshold was 1.25 for V2Rs (**Figure 4-figure supplement 1E**) and H2-M*v* genes (**Figure 4-figure supplement 1F**). This analysis resulted in identification of multiple co-expressing V2Rs and H2-M*v* genes per cell. The chance that a combinatorial pattern identified can be due to a doublet is dependent on the frequency of expression of that gene. Therefore, for abundantly expressed family C V2Rs, it is possible some combinatorial patterns may be from doublets. But in case of family ABD V2Rs, the probability of a particular combination falsely identified due to a doublet more than once is very low. Hence, we assigned a threshold for combinatorial co-expression patterns that are identified in 2 or more cells to be more reliable. These combinatorial expression patterns along with their observed frequency are listed in **supplementary table 7**.

Within the expression patterns represented by a frequency of more than two cells, we observed that family C1 V2R (Vmn2r1) is mostly expressed alone without any other family-C members **(Figure 4F, 4E)** as shown earlier (Silvotti *et al*., 2007). Surprisingly a few cells (31 neurons out of 1025 Gnao1 neurons) were observed to co-express Vmn2r1 along with Vmn2r7 **(Supplementary table 7)**, contrary to earlier reports that Vmn2r1 does not co-express with family-C2 receptors. Most of Gnao1 neurons with family C2 V2Rs (Vmn2r2-2r7) co-express multiple members amongst them, ranging from 2 to 6 V2Rs per cell **(Figure 4F, 4E)**.

In case of family-ABD V2R expression, we observed that most Gnao1 neurons follow ‘one ABD-V2R, one neuron’ rule, except for few that express two ABD-V2Rs **(Figure 4G).** Amongst cells that express more than two ABD-V2Rs, the combination of Vmn2r20 and Vmn2r22 stood out as the highest (44 cells), followed by Vmn2r39, Vmn2r43 and Vmn2r50 (10 cells, **Supplementary table 7**). It is worth noting that these ABD co-expressions in each cell are observed irrespective of whether that cell expresses famly- C1 or C2 V2R **(Figure 4G)**. Among Gnao1 neurons (n1-n4), 229 cells did not express family-C V2Rs **(Figure 4F)** and 46 cells did not express ABD-V2Rs **(Figure 4G)** as per our threshold.

In case of H2-M*v* genes, each cell expresses 1-4 members mostly with famlily-C2 V2Rs **(Figure 4H, 4C, 4D)**. Interestingly, in deviation from this, we observed a pattern whereby H2-M1, -M9 and -M11 genes that show sequence divergence from rest of the H2-M*v* cluster (Ishii, Hirota and Mombaerts, 2003), are specifically expressed in cluster n4 and restricted cells amongst cluster n3, where they co-express with Vmn2r1 GPCR, rather than family-C2 V2Rs (**Figure 4–figure supplement 2B, 2D**). Overall, 56.9% of Gnao1 cells that express either C1-V2R or C2-V2R, did not express any H2-M*v* genes **(Figure 4H)** as per our threshold, reinforcing earlier observations (Ishii and Mombaerts, 2008) that their functional role maybe restricted to a subset of V2R expressing neurons. Furthermore, these H2-M1, H2-M9 or H2-M11 expressing cells along with Vmn2r1, also co-express either Vmn2r81 or Vmn2r82 (V2rf) **(Figure 4–figure supplement 2B, 2C**) as identified before (Ishii, Hirota and Mombaerts, 2003). Two color RNA in situ hybridization with a common probe targeting Vmn2r81 and 82 and separate probes for H2-M1, H2-M9 and H2-M11 shows that Vmn2r81/82 cells always colocalized with the selected H2-M*v* probes **(Figure 4–figure supplement 2E)**.

We performed a similar co-expression analysis on scRNAseq data of p56 VNO from Hills et. al. (Hills *et al*., 2024) and observed that the overall co-expression pattern of V2Rs and H2-Mv genes in Gnao1 neurons matches with ours (**Figure 4-figure supplement 3**). This rules out the possibility of dataset specific artifacts leading to spurious co-expression patterns. Altogether, these data validate the co-expression analysis methodology and identify novel combinatorial co-expression patterns of V2Rs and H2Mv genes.

We next asked, whether the total GPCR or H2-M*v* expression level is regulated or capped in a single cell? To answer this question, we calculated the total family-C-V2R, ABD-V2R and H2-M*v* expression in a single cell that either express one member or multiple members of each family **(Figure 4I-4K)**. The results showed that when cells express more than one V2R, total V2R expression of either family-C or family-ABD (**Figure 4I, 7J)** scales up proportional to total number of expressed V2Rs of that family. This is also true in the case of multiple H2-M*v* expression where total H2-M*v* expression is proportional to the number of H2-M*v* genes expressed in a cell **(Figure 4K)**. As a control, and to eliminate the possibility of doublets, we see that the levels of Gnao1 does not change with number of V2Rs or H2-M*v* genes expressed in a cell **(Figure 4I-4K)**.

Lastly, we performed a similar co-expression analysis for V1Rs with a threshold value of 2.5 (**Figure 4–figure supplement 1D)**. In similarity to the pattern observed for ABD- V2Rs, most Gnai2 neurons expressed a single V1R per cell with some cells co-expressing two V1Rs **(Figure 4–figure supplement 4B, 4A, Supplementary table 8)**. Some of these V1R combinations have been reported recently (Lee, Kume and Holy, 2019). Even in case of Gnai2 neurons, total V1R expression in the cells that co-express two receptors is higher compared to cells that express one GPCR with Gnai2 levels being same **(Figure 4–figure supplement 4C),** indicating that these measurements are from singlet cells. Due to stringent expression threshold, it is possible that VRs expressed at very low level were not considered and thereby leading to neurons where zero VRs are detected (**Figure 4–figure supplement 4B)**. However, these neurons express other genes like neuronal marker Gnai2 at same level as other neurons (**Figure 4–figure supplement 4C)** ruling out the possibility of them being debris. Collectively, these data reveal the patterns of GPCR, H2-M*v* co-expression in VNO neurons, which is likely to be instrumental in deciphering how these neurons respond to single or combination of stimuli, as well as the cellular mechanisms that orchestrate these expression patterns.

### Divergent gene expression profile of mature Gnao1 and Gnai2 neurons

Since exclusion of VR genes did not fundamentally alter the clustering and divergence into mature Gnai2 / Gnao1 neurons, we decided to investigate the gene expression differences between these neurons by performing differential gene expression analysis of mature Gnao1 (cluster 14 from **Figure 1**) and Gnai2 neurons (cluster 7, 16 from **Figure 1**) and identified 924 differentially expressed genes with at least log2 fold change of 1 and adjusted p-value<10e-6. Of these, 456 genes are enriched in Gnao1 neurons and 468 genes in Gnai2 neurons (**Figure 5A, Supplementary Table 9)**. In agreement with previous studies, we see enrichment of V2Rs, H2-M*v*s, B2m (Ishii, Hirota and Mombaerts, 2003; Loconto *et al*., 2003), Robo2 (Prince *et al*., 2009), Tfap2e (Lin, Jennifer M et al., 2018), Calr4 (Dey and Matsunami, 2011) in Gnao1 neurons and V1Rs, Meis2 (Chang & Parrilla, 2016) in Gnai2 neurons (**Figure 5A, Supplementary Table 9**). Along with such known genes, our analysis revealed new differentially expressed genes in the two neuronal subtypes.

**Figure 5.**
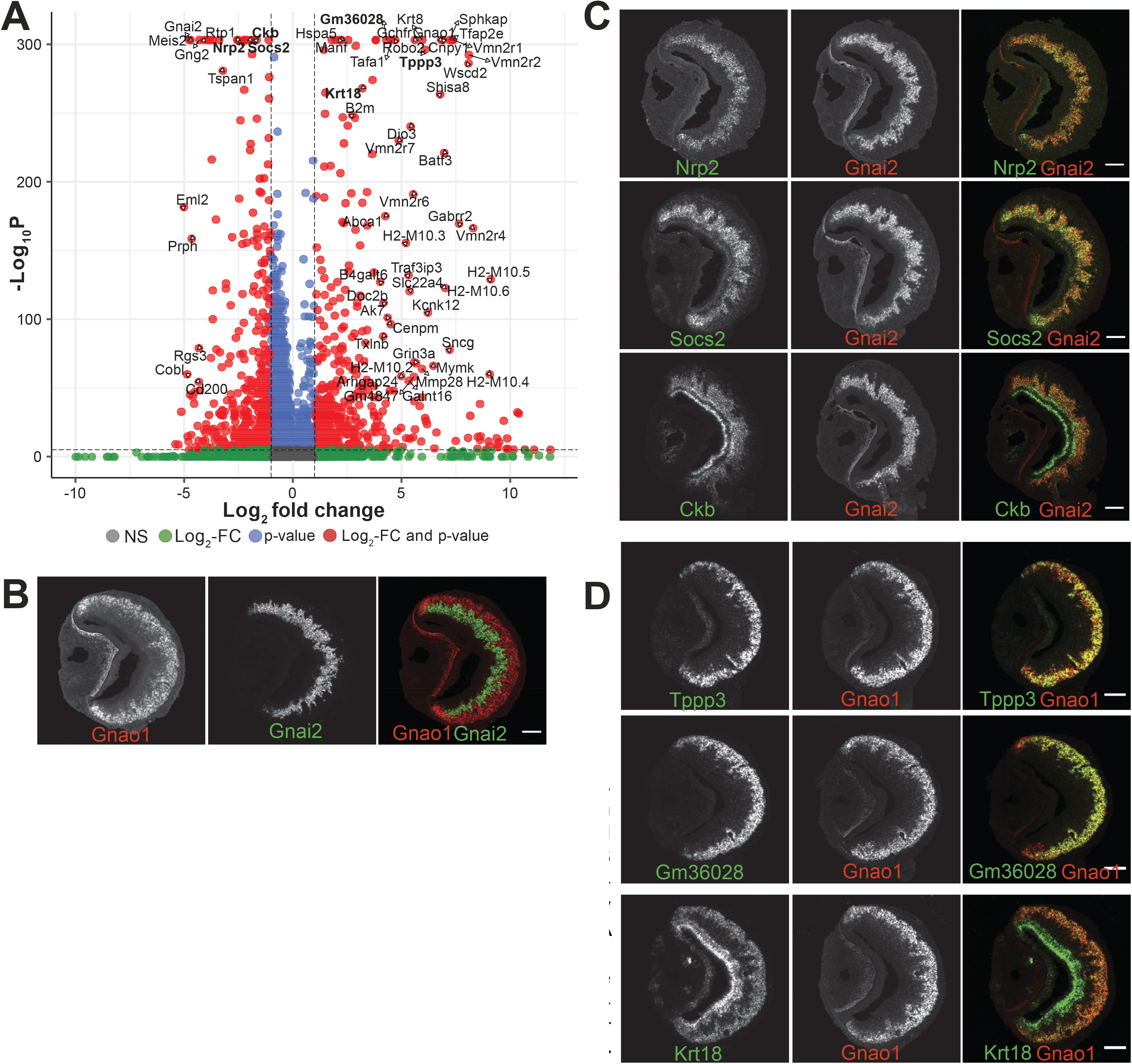
Divergent gene expression: Gnao1 vs Gnai2 neurons. A) Volcano plot showing differentially expressed genes between mature Gnao1 and Gnai2 neurons. Genes that satisfy |log_2_ fold change normalized expression| > 2 (green) and -log_10_ p value > 6 (blue) are considered significant and coloured in red. Non-signifi- cant (NS) genes are colored in grey. Complete list of differentially expressed genes is in **Supplementary table 6**. **C-D)**Two color RNA ISH of selected enriched genes marked in bold on the volcano plot. Gnao1, Gnai2, respective markers of basal and apical neurons are shown in **(B)**. Genes enriched in Gnai2 neurons **(C)** or Gnao1 neu- rons **(D)** are co-localized with the respective markers. RNA-ISH for additional enriched genes are shown in Figure 5**-figure supplement-1**. Scale bar:100 μm.

To confirm differential gene expression amongst mature neurons, we performed two-color fluorescence RNA-ISH of the top candidates from **Figure 5A**, using Gnao1 or Gnai2 as co-labelling markers (**Figure 5B, 5C, 5D)**. Among these genes, Nrp2 and Socs2 expression is restricted to Gnai2 neurons (**Figure 5C**); Tppp3 and Gm36028 expression is restricted to Gnao1 neurons (**Figure 5D)**. Some of the enriched genes such as Ckb in Gnai2 and Krt18 in Gnao1 neurons, were also observed to be expressed in sustentacular cells (**Figure 5C, 5D**). Furthermore, from chromogenic RNA-ISH experiments, we observed genes such as Nsg1, Rtp1, Dner, Qpct, Pcdh7 to be either restricted to or highly expressed in Gnai2 neurons (**Figure 5–figure supplement 1A**) and others such as Apmap, Selenof, Hspa5, Itm2b, Agpat5 were observed to have a gradient of high to low level expression from Gnao1 to Gnai2 zones (**Figure 5–figure supplement 1B**). Two genes: Sncg and Prph specifically expressed in scattered subsets of Gnao1 or Gnai2 neurons respectively (**Figure 5–figure supplement 1A, 1B**) indicating selective expression in a particular neuronal subpopulation. Gene ontology terms associated with the genes validated with RNA-ISH are in **Supplementary table 10**.

These experiments validated that several of the candidate genes identified as differentially expressed from scRNA seq data, are highly specific and spatially localized to one of the neuronal sub-populations. A complete list of differentially expressed genes is in **Supplementary table 9**. Taken together our data indicate that VNO neurons that start from a common progenitor, go on to transiently express genes during immature stages that could decide their cell fate and these cells exhibit markedly different gene expression profiles in the mature stage.

### Differential ER environment in Gnao1 and Gnai2 neurons

Given that Gnao1 and Gnai2 neurons express divergent GPCR families, we asked the question whether the differentially expressed genes indicate fundamental differences in cellular or molecular processes between these two cell types. We performed gene ontology (GO) enrichment analysis on the two sets of genes. Interestingly, we observed that amongst the Gnao1 neuron genes, most of the enriched terms with p value <0.05, are related to endoplasmic reticulum (ER) processes, which include ‘protein folding’, ‘response to ER stress’ and ‘ERAD pathway’ (**Figure 6A**). Conversely, a gene ontology analysis of Gnai2 neuron genes did not reveal any particular enrichment of GO terms in this neuronal subset. As more than 20% of the genes associated with protein folding **(Figure 6A)** are enriched in Gnao1 neurons, we sought to probe the spatial transcription pattern of some of the candidates. Fluorescence RNA in-situ hybridization confirmed the selective expression of ER genes such as Creld2, Dnajc3, Pdia6 and Sdf2l1 amongst Gnao1 neurons (**Figure 6B**).

**Figure 6.**
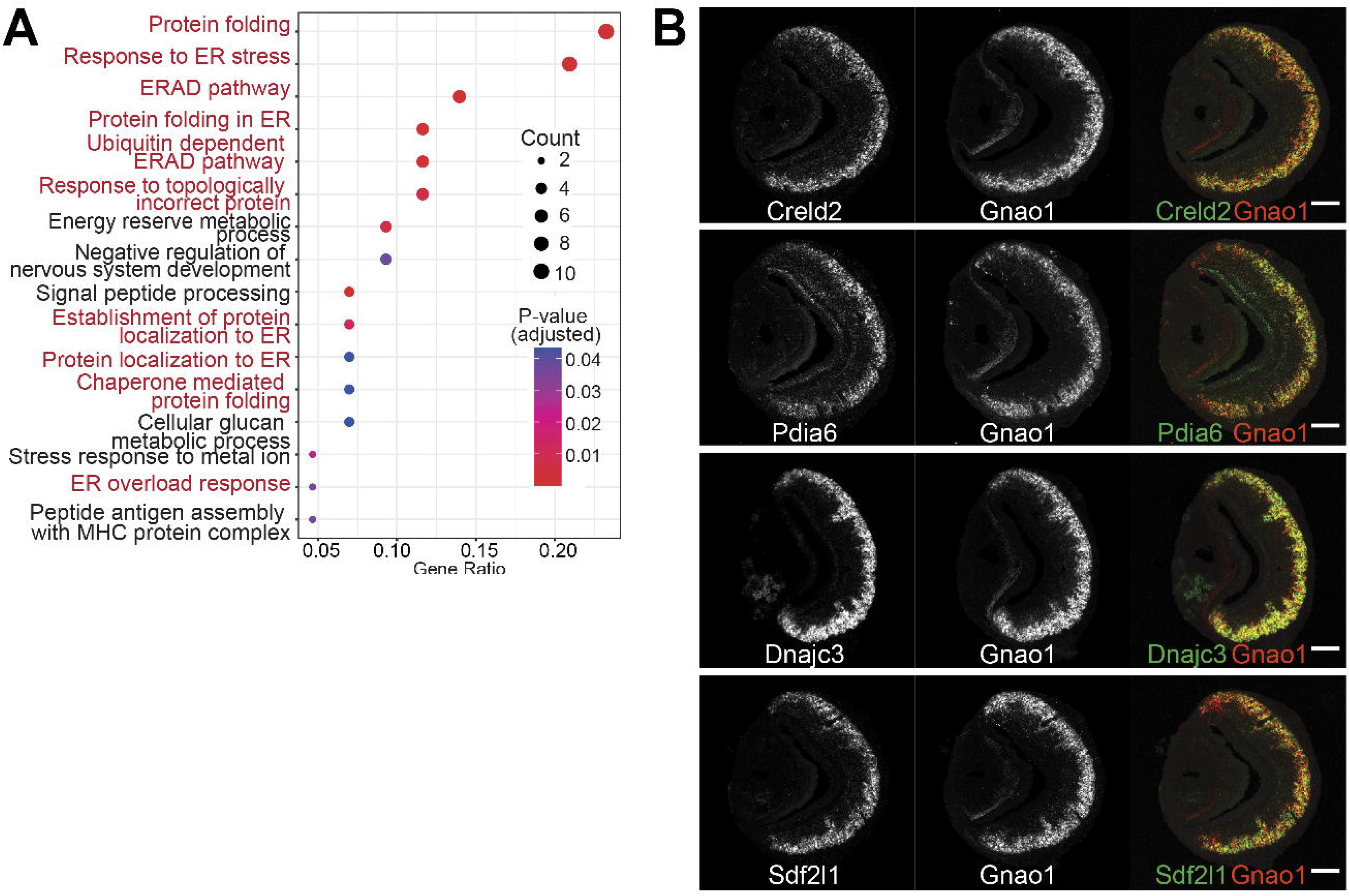
A) ER gene expression in Gnao1 neurons. Annotation of gene ontology (GO) biological processes of genes that are significantly enriched in Gnao1 neurons from Figure 5A. GO terms related to ER processes are marked in red. p-value < 0.05 was used as cut-off. **B)** VNO coronal section two color fluorescent RNA-ISH of selected Gnao1 enriched ER chaperone genes (Creld2, Pdia6, Dnajc3, Sdf2I1: green), co-labelled with Gnao1 probe (red) shows Gnao1 zone restricted expression of these genes. Scale bar 100µm.

To test whether these findings extend to the protein level, we performed immuno- fluorescence microscopy of VNO sections using antibodies specific to a collection of ER proteins. We first tried an antibody against SEKDEL, a common ER retention signal, which intensely labelled Gnao1 neurons (**Figure 7A**). This selective labelling was confirmed by co-labelling with anti-Gnao1 (**Figure 7B, 7D**), where anti-SEKDEL ER signal is seen in Gnao1 neurons. We quantified enrichment of SEKDEL across multiple (26) VNO sections from 3 mice, labelled with anti-SEKDEL, anti-Gnao1 and anti-Omp as a control. Normalized fluorescence intensity quantified along the apical-basal axis (**Figure 7D, 7E**), shows that SEKDEL signal intensity increases along with Gnao1. Surprisingly, several other antibodies that detect ER chaperone proteins: Hspa5 (BiP), PDI, Grp94, Calnexin (**Figure 7F**, **Figure 7-figure supplement 1A**); as well as antibodies against ER membrane / structural proteins: Sec61β, Atlastin1, NogoB, Climp63, Reep5 (Goyal and Blackstone, 2013) show a similar enrichment in Gnao1 neurons, as seen from the immune-fluorescence images and their quantification with Gnao1 or SEKDEL antibodies (**Figure 7G,H**, **Figure 7-figure supplement 1B**). Some ER protein localizations (SEKDEL, Hspa5, PDI, Atlastin1, NogoB) were detected selectively amongst basal zone Gnao1 neurons, while others (Sec61β, Calnexin, Grp94, Reep5, Climp63) detected a pattern of Gnao1 neuron enrichment, with lower levels in Gnai2 neurons. To test whether higher levels of these ER proteins resulted from their increased transcription, we compared their RNA levels in Gnai2 / Gnao1 neurons from our scRNA dataset (**Figure 7–figure supplement 2**). In case of Hspa5 (Bip), Hsp90b1 (Grp94), Sec61b mRNA levels are higher in Gnao1 neurons (also see Hspa5 in **Figure 5A**, and RNA-ISH in **Figure 5-figure supplement 1**). On the other hand, RNA level of Atlastin1, PDI were comparable (**Figure 7–figure supplement 2**), indicating that Gnao1 neuron enrichment of some ER proteins could also arise due to post-transcriptional regulatory mechanisms.

**Figure 7.**
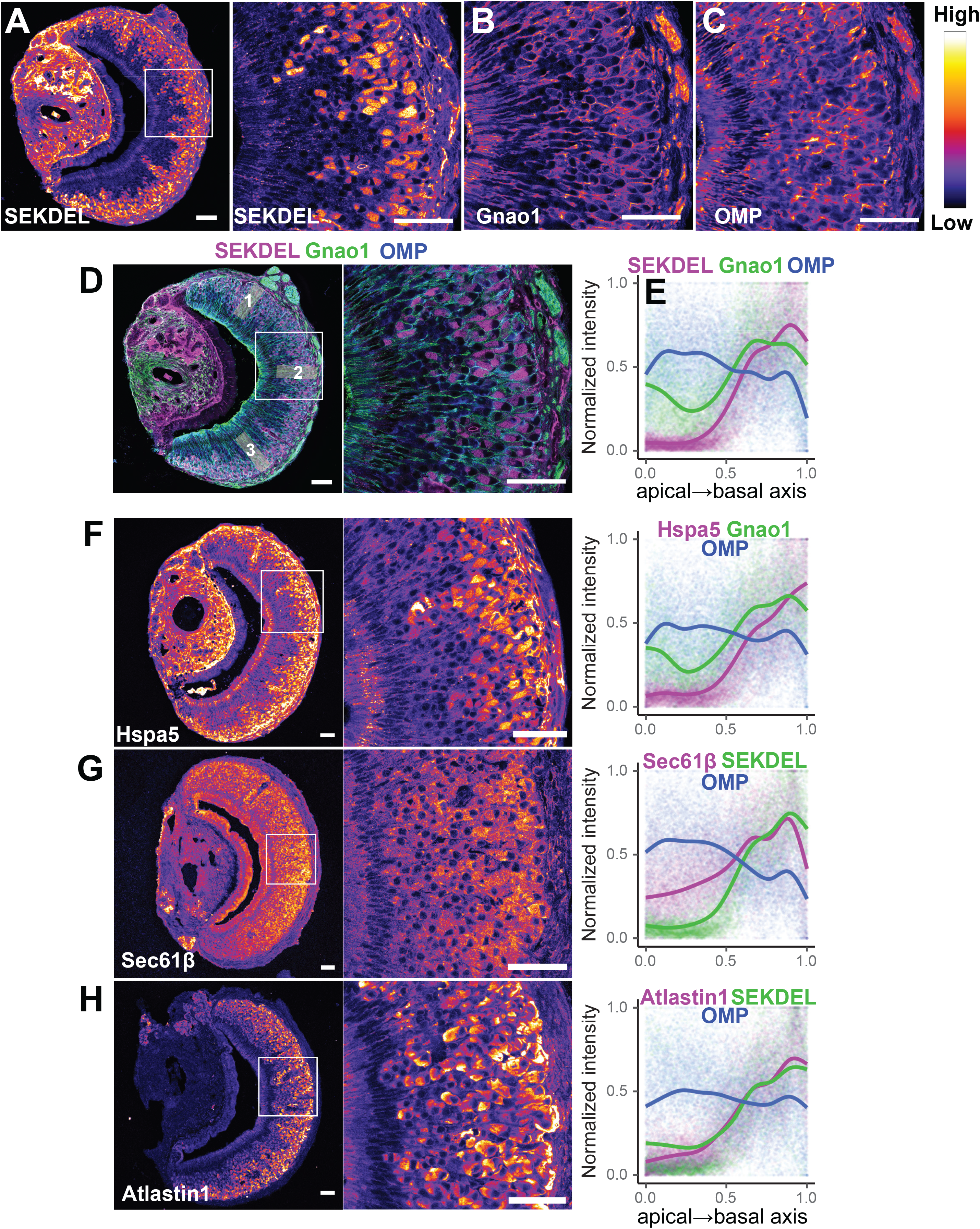
**Differential localization of ER proteins in VNO neurons.** Pseudocolored immunofluorescence images of VNO coronal sections labelled with antibodies against KDEL (anti-SEKDEL), a common ER retention signal **(A)**. The section is co-labelled with anti-Gnao1 to mark basal zone neurons **(B)** and anti-OMP to mark all mature neurons **(C).** Signal intensity of KDEL, Gnao1, OMP channels is quantified from ROIs along the apical-basal axis as shown in example **(D)**. Signal intensity measured from multiple sections (n>20 for each anitbody) from 3 animal replicates was normalized and trendline was fitted to show the Gnao1 neuron enriched localization of anti-KDEL **(E)**. Points on the plot show normalized intensity values color coded for each antibody on which the trendline was fitted. Similar imuno-labelling and quantification of ER chaperone Hspa5 (BiP) **(F)**, ER membrane WUDQVRORFRQ VXEXQLW 6HF61ȕ **(G)** and ER membrane protein Atlastin1 **(H)** indicate their enrichement in Gnao1 neurons compared to Gnai2 neurons. Distribution of additional ER chaperone and membrane proteins is shown in Figure 7**-figure supplement-1**. Scale bar: 50 μm.

Many of the Gnao1 enriched ER genes and their protein products are chaperones, while others are ER structural proteins. The upregulation of both types of ER proteins prompted us to examine the cells by electron microscopy. Serial sections (70 nm thick) of ∼1 mm square regions of the VNO were collected on tape and were imaged by scanning electron microscopy (Baena *et al*., 2019). Since the sections contained the entire VNO, we were able to distinguish Gnai2 from Gnao1 cells by their location as they are spatially segregated into apical or basal zones respectively within the neuroepithelium (**Figure 8A**). In Gnao1 cells, more than half of the cytoplasm contained dense smooth ER with cubic membrane morphology (Almsherqi, Kohlwein and Deng, 2006; Almsherqi *et al*., 2009), also known as organized smooth ER (OSER) (Snapp *et al*., 2003) (**Figure 8A, 8B,** **Figure 8–figure supplement 1**). In remainder of the cytoplasm, there were closely apposed parallel sheets of ER membrane that were contiguous with lamellar ER surrounding the nucleus (**Figure 8–figure supplement 1**). The apical/Gnai2 neuronal cell bodies are smaller than those of Gnao1 cell bodies. They also appeared to have cubic ER membranes, but these seemed denser and occupied a smaller fraction of cytoplasmic volume (**Figure 8A, 8C**).

**Figure 8.**
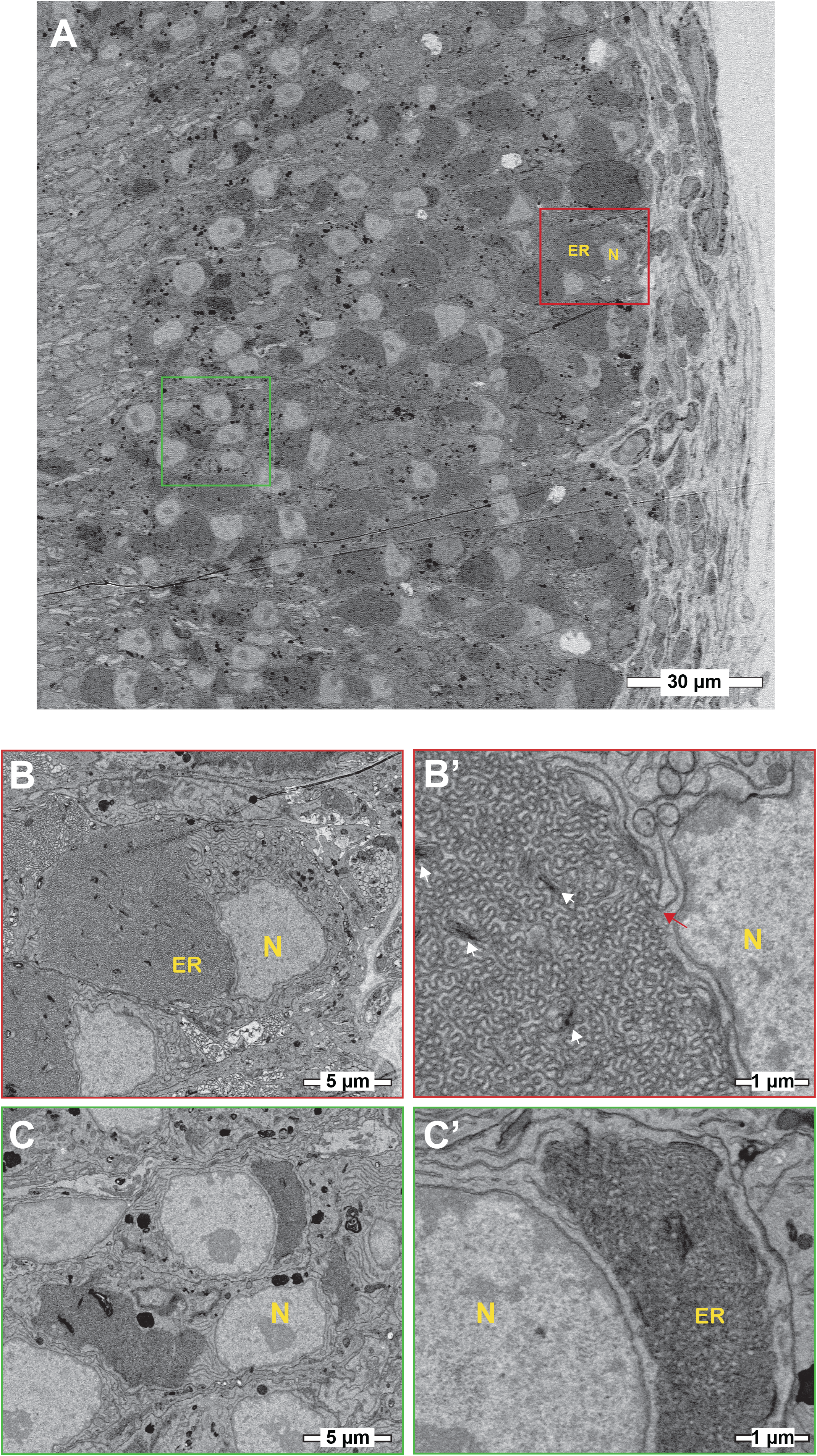
Basal/Gnao1 neurons are densely packed with cubic membrane ER. A) Scanning electron micrographs of vomeronasal sensory epithelium at low magnification showing cell bodies of VSNs, sustentacular cells and basal lamina. Boxed regions in red or green mark cell bodies of basal/Gnao1 or apical/Gnai2 neurons respectively, that are displayed at higher magnifica- tion below. Nucleus (N) appears light and ER is dark. Cell bodies of basal/Gnao1 neurons are larger and occupied by substantial amount of ER, incomparison to apical/Gnai2 neurons. **B, B’)** Magnified micrographs show the cell body of a basal/Gnao1 neuron, packed with cubic ER membranes. White arrows point to dense membrane stacks that are better resolved in Figure 8**-figure supplement 1**. Red arrow points to lamellar ER membrane that is contiguous with the cubic membrane. **C, C’)** Cell bodies of apical/Gnai2 neurons also seem to show dense cubic membrane ER, that is smaller in comparison to basal/Gnao1 neurons.

Taken together, our data indicates that mature Gnao1 neurons differ substantially from Gnai2 neurons in selectively upregulating the transcription of several ER genes. Furthermore, this neuronal subset also exhibits higher levels of ER proteins, via transcriptional and post-transcriptional mechanisms, as well as a hypertrophic smooth ER that is arranged in the form of organized cubic membrane ultrastructure. Collectively, these observations indicate a specialized ER environment that could play a significant functional role in the Gnao1 subset of VNO neurons.

## Discussion

The vomeronasal organ has been a model of considerable interest for molecular sensory biology, to study the diversification of sensory neurons, uniquely evolved gene families and their patterns of expression, all of which are essential to understand how specific social chemo-signals elicit innate behaviors. In this study, using single cell and low-level spatial transcriptomics, we developed a comprehensive resource identifying cell types of the VNO neuroepithelium and studied the developmental and receptor co-expression patterns within sensory neurons. We organized our dataset as an interactive web portal resource, accessible from www.scvnoexplorer.com, where a gene can be queried for its expression pattern and levels amongst cell clusters.

The identification of distinct cell clusters and validation of marker expression through RNA *in situ* hybridization, revealed a heterogeneous cellular composition of the vomeronasal neuroepithelium. In particular, we identified specific new gene markers, such as Ppic in sustentacular cells and Ly6d in non-sensory epithelium, which would be helpful in future to genetically tag or target these cell types and to study their developmental origins. Our study also identified specific types of macrophage cells within the neuroepithelium. Given the increasing appreciation of the role of tissue resident or infiltrating macrophages in a variety of physiological processes (Nobs and Kopf, 2021; Mass *et al*., 2023) it is tempting to speculate that macrophages could also play an important role in VNO neuronal differentiation or the regulation of inflammatory/immune responses to external insults. Future experiments, performing targeted purification, deep sequencing, and functional manipulation of VNO macrophage types could provide deeper insights on their role in VNO function and whether they share cellular identity with MOE macrophages.

### Development of VSNs

VNO sensory neurons expressing Gnao1 and Gnai2 develop continuously throughout life from a single progenitor cell population located within the VNO marginal zones and differentiate to immature and mature neurons. Recent developmental studies have demonstrated that the dichotomy is established by Notch signaling followed by transcription factor, Bcl11b expression that marks the progenitor cells to take Gnao1 fate (Enomoto *et al*., 2011; Katreddi *et al*., 2022). Most Importantly, a critical period of Notch signaling determines whether progenitors give rise to apical, basal or sustentacular cell types. After Gnao1/Gnai2 identity is established, transcription factor Tfap2e, continuously expressed from immature to mature Gnao1 neurons, is required to maintain the Gnao1 fate (Lin *et al*., 2018, 2022; Katreddi *et al*., 2022). Pseudo-time analysis on our data confirmed a developmental trajectory that starts with progenitor cells, leading to immature neurons that take on either Gnai2 or Gnao1 fate. From our scRNA seq data, a comparison of immature neurons revealed that apart from Tfap2e, transcription factors: Creb5, Prrxl1, Lmo4, Foxo1 also express in immature Gnao1 neurons. But unlike Tfap2e, Creb5 and Prrxl1 expression is restricted to only immature stage of Gnao1 neurons. Although, Foxo1 and Lmo4 continue to express in mature Gnao1 neurons, their RNA levels are high in immature Gnao1 neurons and reduce upon maturation. Among the transcription factors identified, Prrxl1 has been shown to involved in the development of dorsal root ganglion neurons where it has been shown to autoregulate its expression (Monteiro *et al*., 2014). Our observations on the tight temporal regulation of these transcription factor related genes in the developmental trajectory of Gnao1 neurons indicates that additional molecular players might be important in further specification or maintenance of this neuronal lineage. Further experiments are needed to tease out the mechanism and precise choreography by which these transcription factors collectively specify and maintain the Gnao1 cell identity as well as gene expression patterns unique to these neurons.

### Receptor co-expression in Gnao1 neurons

One of the key features distinguishing basal zone Gnao1 VNO neurons from apical Gnai2 or MOE neurons, is the combinatorial co-expression of two or more V2R family GPCRs along with H2-M*v* class-1b MHC molecules in each cell. Our analysis of receptor expression in mature VNO neurons, revealed that members of V1R and V2R-ABD gene families largely follow the one-neuron-one receptor rule, but with multiple exceptions. Some V1Rs were found to consistently co-express in Gnai2 neurons and the major combinations observed from our analysis (such as Vmn1r85/86, **Figure 4–figure supplement 4A**), match those identified recently from a functional single cell study (Lee, Kume and Holy, 2019), leading confidence to the methodology used in our co-expression analysis. Receptors of the ABD-V2R subfamilies are also mostly expressed at one receptor per neuron, with notable exceptions being the observed combinations of Vmn2r20/22 and Vmn2r39/43/50 being co-expressed per cell.

In our study, the analysis of family-C, family ABD-V2Rs and H2-M*v* genes expressed per cell, confirmed the earlier reported sub-division of basal/Gnao1 neurons into those that express either Vmn2r1 (family-C1) or Vmn2r2-Vmn2r7 (family-C2), resulting in a broad four-part division of mature VNO neurons: those expressing a) Gnai2/V1Rs, b) Gnao1/Vmn2r1, c) Gnao1/Vmn2r2-2r7; H2-M*v*+, d) Gnao1/Vmn2r2-2r7; H2-M*v*-. The co-expression of ABD-V2R sub-families A1-A5 and most of the H2-M*v* genes with Vmn2r2/family-C2 neurons **(Figure 4C-E, cluster n3, n2, Supplementary table-7)** as well as preferential co-expression of A6-A9, B, D, E sub-families with Vmn2r1 along with exclusion of H2M*v* genes from these neurons **(Figure 4E)**, points to the overall non-random logic of V2R/H2Mv co-expression. It would be interesting to see how receptor de-orphanization and future functional experiments will map onto these elaborate co-expression patterns. At the same time there were notable deviations observed from these rules, namely the restricted and sparse expression of H2-M1/M9/M11 with Vmn2r1 neurons in cluster n4 (**Figure 4E**, **Figure 4–figure supplement 2B, 2E)**, where the receptors Vmn2r81/82 are also expressed selectively in these neurons.

Of note, the combined expression of VR/H2-M*v* in a cell is not capped and is proportional to number of VRs and H2-Mv genes expressed in that cell (**Figure 4I-4K**). It would be interesting to see whether the levels of co-expressing transcripts also translate to protein levels in cells and their impact on functional interactions, if any, between co-expressed proteins.

### ER environment in VNO neurons

When comparing gene expression amongst Gnai2/Gnao1 mature neurons, what stood out was the unexpected enrichment of gene ontology terms associated with ER functions within Gnao1 neurons **(Figure 6A, 6E)**. Most of these ER-Gnao1 enriched genes from our dataset, match the ones identified from a similar comparison by Lin et al 2022, supporting our findings. We validated several of these ER genes via RNA-ISH to be exclusively expressed or enriched in Gnao1 neurons. Even more puzzling was our observation that this enrichment pattern extended at the protein level to many ER proteins probed by immune labelling (**Figure 7**). In some cases, both RNA and proteins were Gnao1 enriched, whereas in others, RNA levels were comparable but protein levels were biased towards Gnao1 cells. The enrichment of ER genes/proteins via transcriptional and/or post-transcriptional mechanisms presents a new feature of these cell types that could be associated with their differentiation and function. High levels of several chaperones such as Creld2, Dnajc3, Pdia6, PDI, Hspa5/Bip, Grp94, Calnexin in Gnao1 neurons **(Figure 6B**, **Figure 7F**, **Figure 7-figure supplement 1A)** compared to Gnai2 neurons indicates a chaperone rich ER environment in these neurons. The chaperone Calr4 has been shown to be enriched in Gnao1 neurons (Dey and Matsunami, 2011). Our observations indicate that the enrichment of chaperones is a generalized feature of Gnao1 neurons that extends to several other genes/proteins of related function and localization.

Since it has been proposed that the V2R family is an evolutionarily recent acquisition in rodents and marsupials (Takigami *et al*., 2004) and sub-family members, such as sub family-C2, A1-A6 V2Rs, as well as H2-M*v* genes have evolved in Muridae (Young and Trask, 2007; Silvotti *et al*., 2011), it is possible that mouse Gnao1 neurons may have required to co-evolve ER protein folding mechanisms to handle multiple GPCR/H2-M*v* co-expression and their putative protein interactions. In addition to our observations on the generalized upregulation of ER genes in Gnao1 neurons, it may be possible that some specific ER genes are also necessary for proper folding of vomeronasal type-2 GCPRs. These data and observations assume significance given the well-recognized fact that functional reconstitution of these GPCRs and H2-M*v* molecules in heterologous cells remains a persistent challenge, with some success being reported using the chaperone Calr4 (Dey and Matsunami, 2011). On the other hand, olfactory receptors have been shown to co-opt the ER unfolded protein response (UPR) pathway via the ER proteins PERK/CHOP and transcription factor ATF5 as a mechanism for their functional expression, setting the stage for ONOR expression pattern (Dalton, Lyons and Lomvardas, 2013). ATF5 is observed in the VNO, however, it is reported to be broadly expressed across Gnai2 and Gnao1 neurons (Nakano *et al*., 2016; Dalton, 2018; Dalton *et al*., 2018),. Further experiments are required to evaluate whether V2Rs and Gnao1 neurons adopt similar mechanisms and how the chaperone rich ER environment observed here might impact the expression, folding of GPCRs and neuronal function.

It is worth noting that in addition to the expression of ER chaperones in Gnao1 neurons, we also observed distinctly higher levels of ER structural and membrane proteins. Levels of membrane curvature stabilizing proteins - Reep5, NogoB/Reticulon4B, the three-way junction formation protein - Atlastin1 (Goyal and Blackstone, 2013), ER sheet lumen spacer protein - Climp-63/Ckap4 as well as Sec61β - a commonly used ER membrane marker were all observed to be high in Gnao1 neurons **(Figure 7G, 7H,** **Figure 7-figure supplement 1B).** Thus, it is possible that an added layer of complexity could involve modulation of total ER content or ER structure in Gnao1 neuronal subtypes. Our electron microcopy results **(Figure 8**, **Figure 8-figure supplement 1)** revealed a strikingly higher smooth ER membrane content in Gnao1 neurons compared to Gnai2 neurons. Interestingly, this ER adopts a cubic membrane architecture, packing the Gnao1 neuron cell body with gyroid / sinusoidal membrane. Early electron microscopy studies in rodents, identified that smooth ER content is higher in VSNs than olfactory sensory neurons (Ciges *et al*., 1977; Breipohl *et al*., 1981; Taniguchi and Mochizuki, 1982), however, these studies preceded the discovery of Gnao1-Gnai2 dichotomy. To our knowledge, Breipohl and colleagues (Breipohl *et al*., 1981) reported the presence of smooth ER whorls in VSNs, later termed as cubic membranes. Future studies using serial section EM would be instrumental in reconstructing the 3D architecture of VSN ER.

Cubic membranes, representing highly curved periodic structures, have been observed in a variety of biological contexts, however their functional significance has been harder to pin down (Almsherqi, Kohlwein and Deng, 2006; Almsherqi *et al*., 2009). In the context of VNO, and in particular Gnao1 neurons, we speculate that the high ER content and dense packing in the form of cubic membranes, could indicate a perpetual stress like state in these neurons, necessary to address unique folding requirements of V2R or H2- M*v* proteins. For instance, maturation of these proteins and their proper folding or multimerization may require a slower transit through the ER, necessitating the sinusoidal arrangement of ER membranes. Likewise, the V2R biosynthetic processes in Gnao1 neurons may require enhanced levels of ER chaperones, which could in turn induce ER membranes into a homeostatic response. Can V2Rs themselves trigger upregulation of ER chaperones and the hypertrophic ER, or are these cellular characteristics established before the onset of V2R expression? From our single cell transcriptomics data, we tried to identify at what stage ER chaperones express during the developmental trajectory of VSNs. We observed that onset of Vmn2r1 expression coincides with upregulation of ER chaperones in cluster n6 immature Gnao1 neurons (**Figure 9** **A, B)**. However, some Gnao1 cells show upregulated ER gene expression without Vmn2r1/Vmn2r2 expression (**Figure 9B**), hence further experiments are needed to dissect the mechanism and precise relationship between neuronal differentiation, GPCR expression and ER chaperones as well as ER ultrastructure.

**Figure 9.**
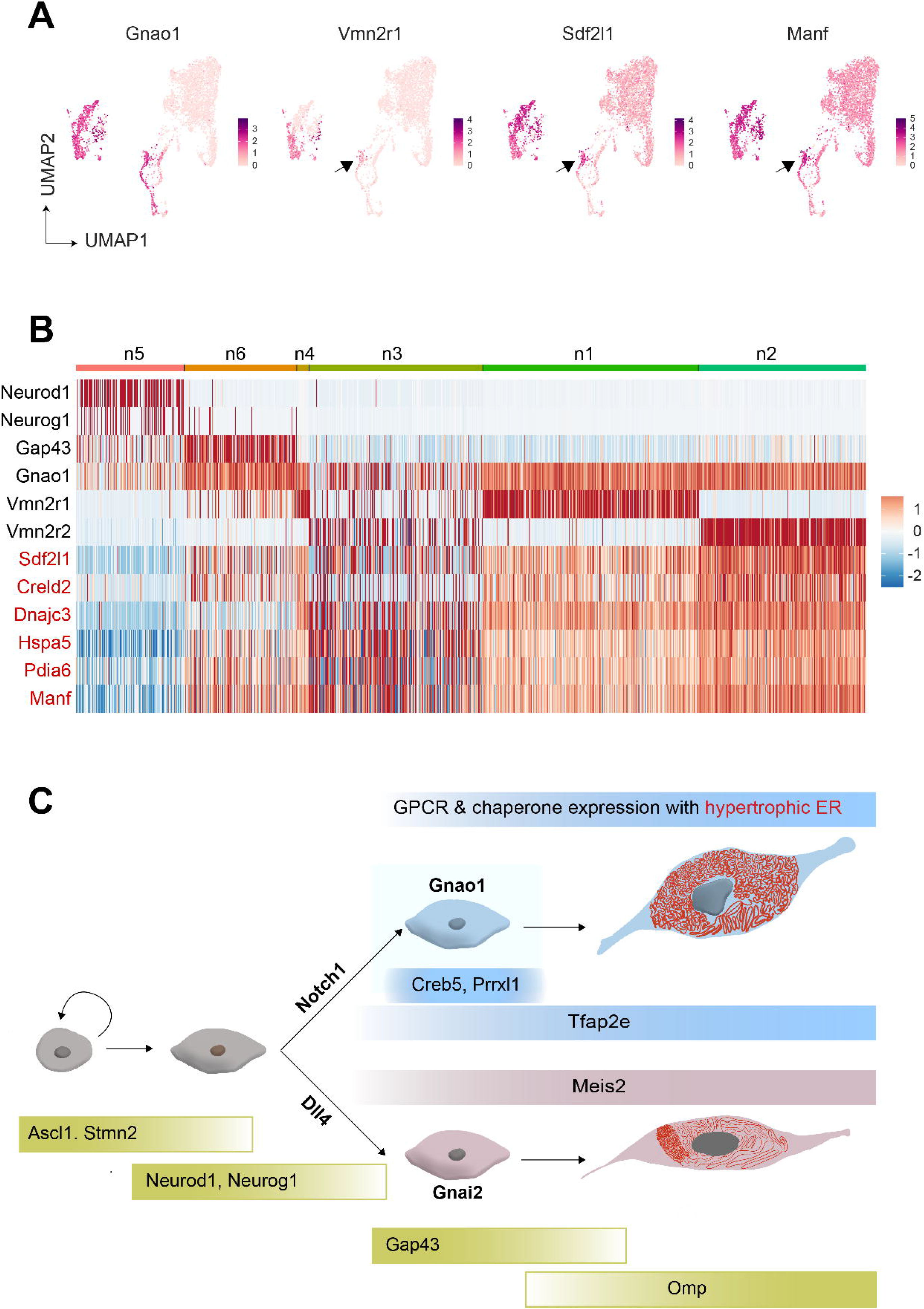
Onset of V2R expression coincides with the expression of ER chaperone genes. A) Feature Plot showing the expression of Gnao1, Vmn2r1, Sdf1I1 and Mani. Sdf2I1 and Mani are known **ER chaperones and their upregulation in Gnao1 neurons coincides with Vmn2r1 expression, which is** preceded by Gnao1 expression. **B)** Heatmap showing the expression of Gnao1, Vmn2r1, Vmn2r2 and several ER chaperone genes (red) in the clusters arranged as per their developmental trajectory. **C)** Cartoon **summarizing major transcription factor expression during development leading to Gnao1 neurons with** chaprone rich hypertropic ER compared to Gnai2 neurons.

Overall, our data opens a new aspect to look at Gnao1 neurons, when understanding their function: a highly specialized ER environment that is divergent from Gnai2 neurons. The ER genes and their Gnao1 biased expression we identified, could be downstream targets of specific transcription factors and as a result of neuronal differentiation, while functionally these could be required for the proper folding and co-expression of V2R GPCRs. The developmental trajectory of VSNs into Gnai2/Gnao1 neurons via transcription factors and their differential ER environment is summarized as a model in **Figure 9C**.

In conclusion, the comprehensive scRNA seq analysis presented in this study contributes valuable insights into the complexity of the vomeronasal neuroepithelium, offering a roadmap for further investigations into the molecular mechanisms underlying sensory perception and neural development in this specialized sensory organ. Vomeronasal neurons may also serve as model system to study specialized ER structure-function relationship and its role in maturation of GPCR expressing neurons.

## Materials and Methods

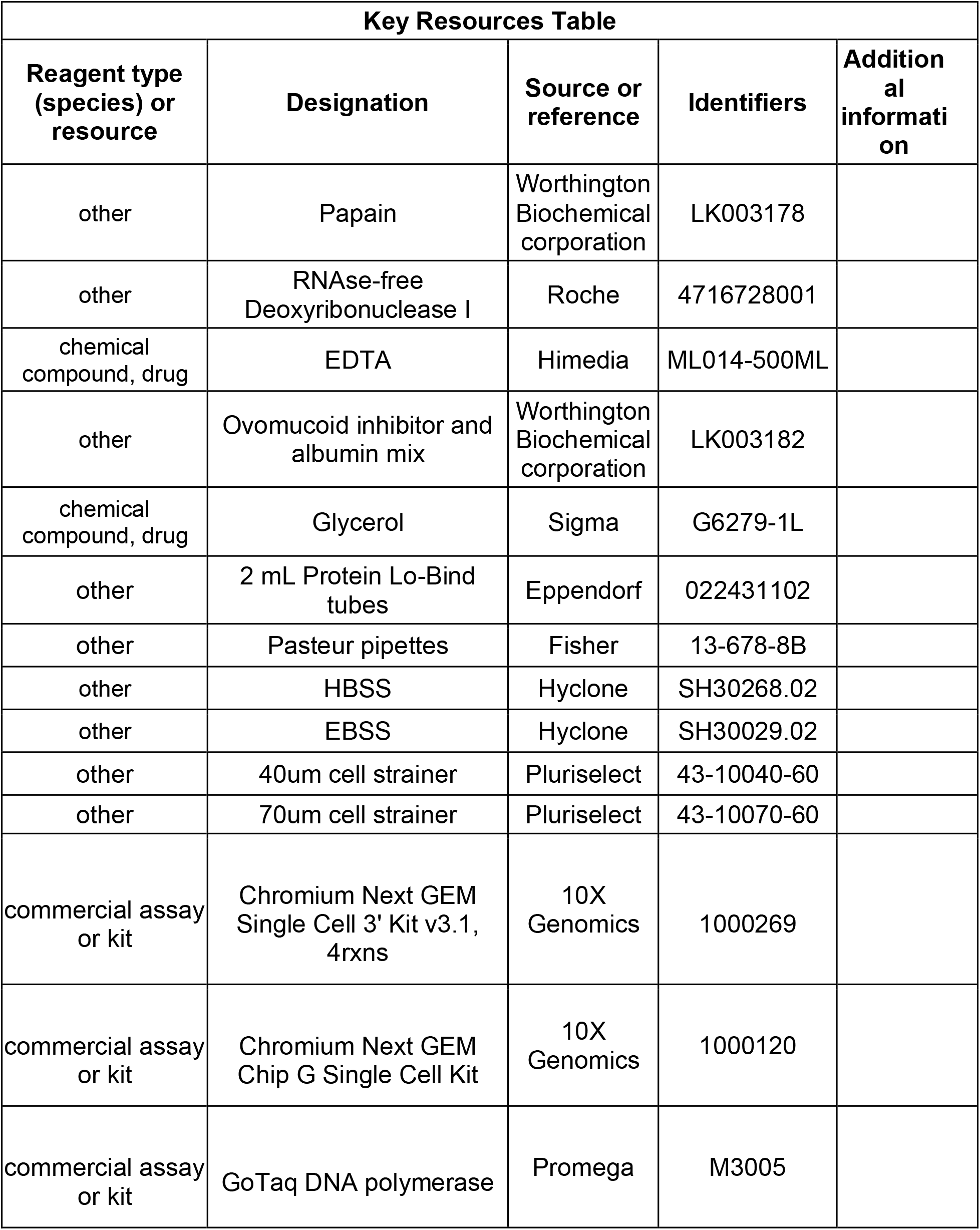
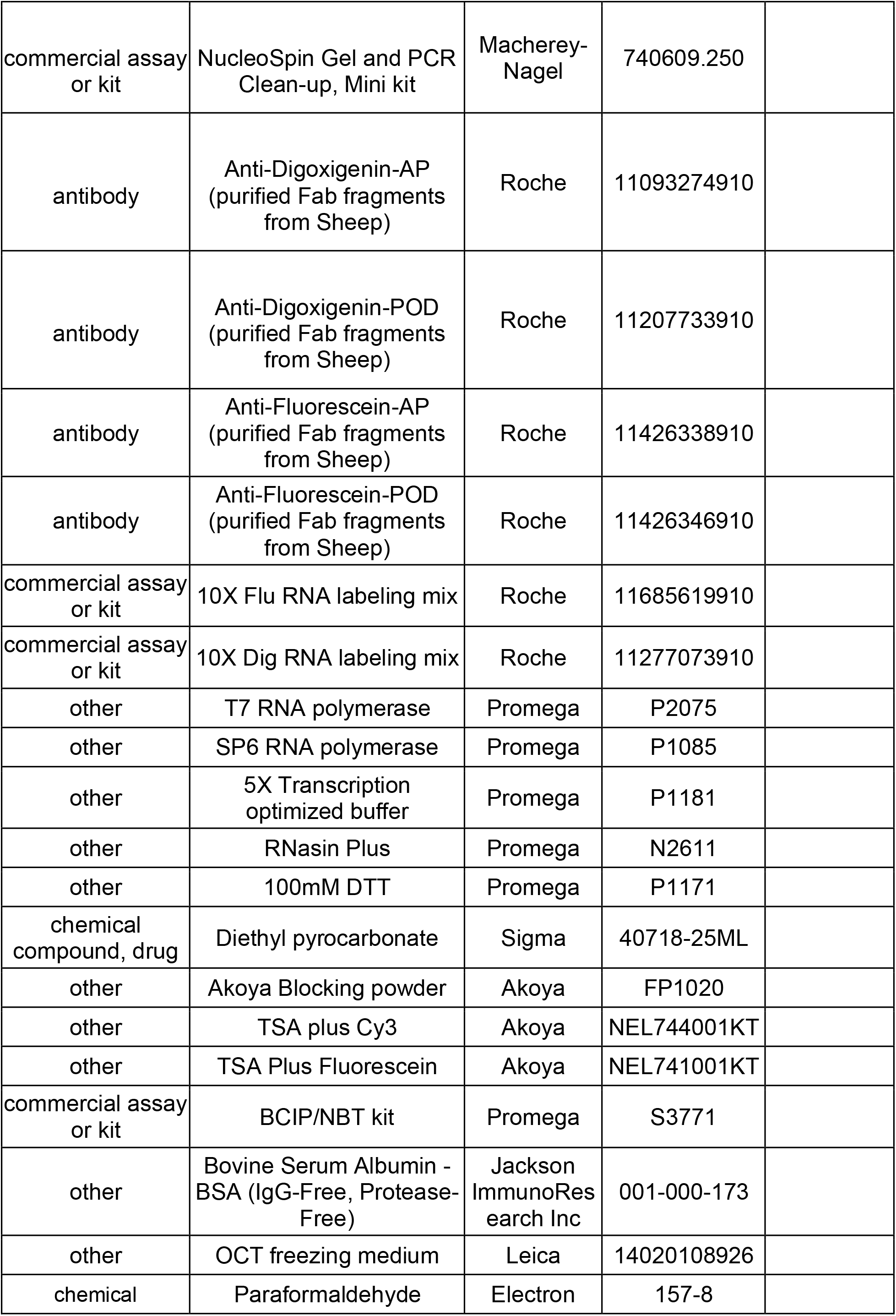
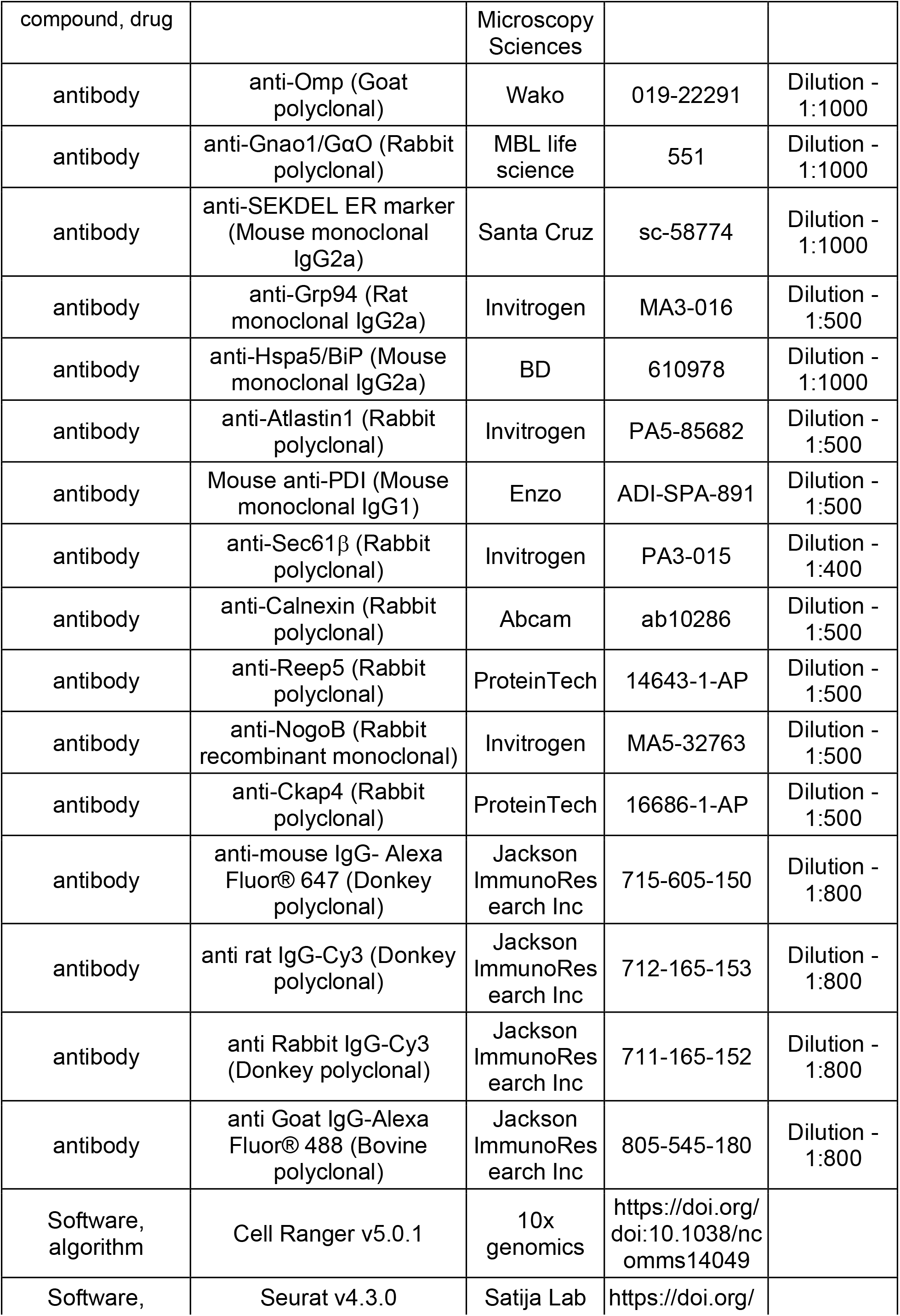
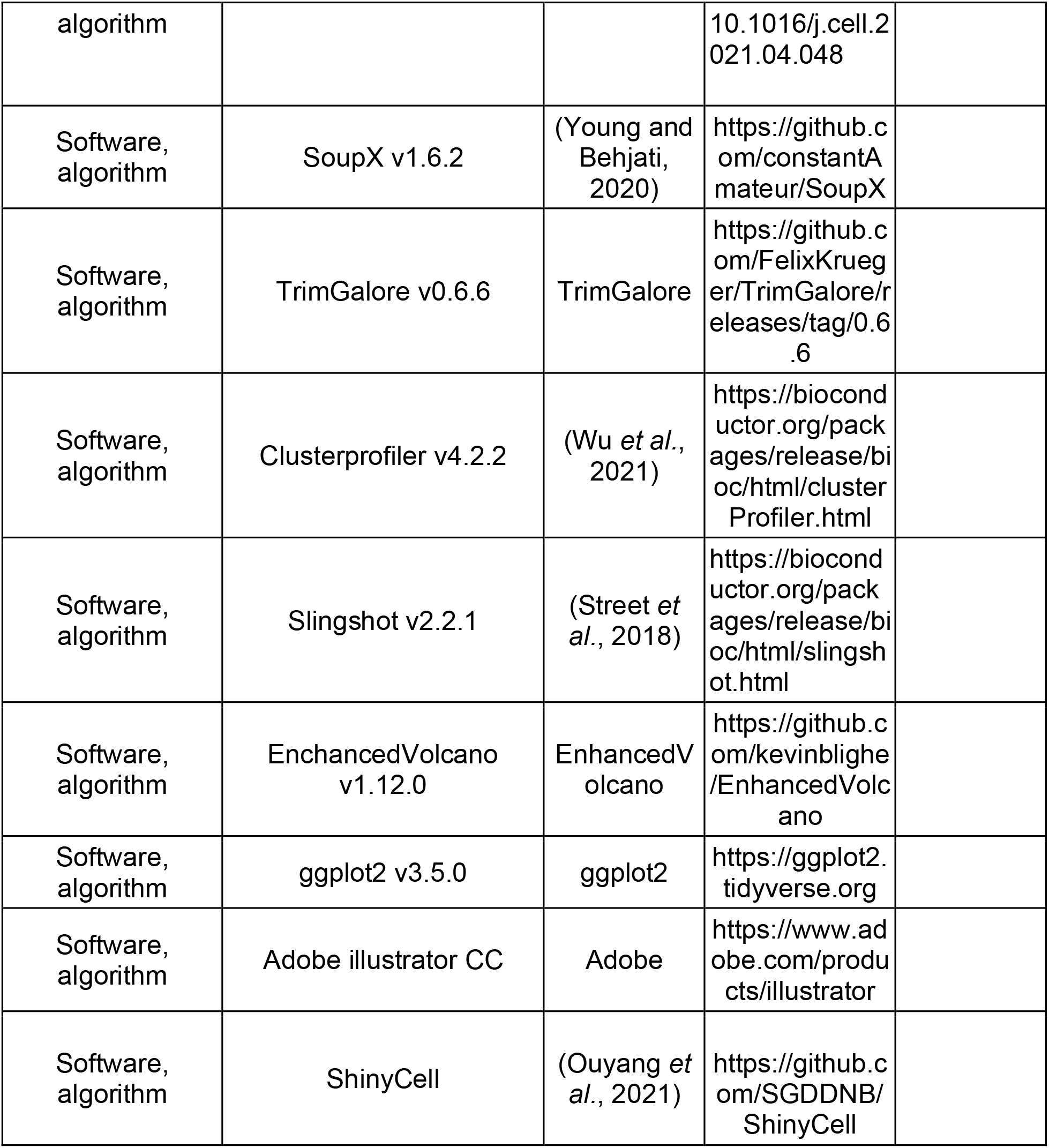

### Animals

C57BL/6J mice were purchased from JAX (Strain #000664) and were used for all experiments. Mice were housed in a specific pathogen-free barrier facility, with a 12-hour light-dark cycle and ad-libitum provision of feed and water. For single-cell RNA sequencing experiments, male and female mice were weaned at postnatal 3 weeks age, followed by housing in separate individually ventilated cage racks to avoid exposure to opposite sex stimuli, and used at 7-8 weeks age. All experiments were carried out with approval from the Institutional Animal Ethics Committee of TIFR Hyderabad.

### VNO dissociation

Papain dissociation buffer (PDB) was made by reconstituting single use papain vial from Worthington Biochemical Corporation with 5 mL Earle’s balanced salt solution (EBSS) (pH 7.2) and warmed to 37°C until solution appears clear by maintaining 95% CO2, 5% O2 environment. VNOs were dissected from male and female animals (6 male, 10 female) and processed separately. The sensory epithelium was separated using forceps and immediately placed in EBSS equilibrated with 95% CO2, 5% O2. 60U of DNase was added to the prewarmed PDB made earlier and 3mL of it is transferred to a single well of 12- well dish. Neuroepithelial tissue from multiple animals was transferred to this well containing the final dissociation buffer and was cut into small pieces and the suspension was transferred to 14 mL tube and was incubated with gentle shaking at 37°C for 30 minutes on a thermomixer (Eppendorf) by passing 95% CO2, 5% O2 on top of the liquid headspace. During this incubation, the solution was triturated with fire-polished Pasteur pipette every 10 minutes and at the end the incubation. After incubation, the suspension was passed sequentially through 70µm, 35µm filters to remove tissue debris. The filtered suspension was further layered over a mix of ovomucoid inhibitor (2.5 mg/mL) and albumin (2.5 mg/mL) in EBSS and centrifuged at 400 x g for 5 minutes at room temperature to remove subcellular debris or membrane fragments. The supernatant was discarded, and the cell pellet was used for next steps after Hank’s balanced Salt solution (HBSS) wash.

### Single-cell library preparation and sequencing

Dissociated neurons resuspended in HBSS were used for library preparation. Using hemocytometer and trypan blue staining, cell density was estimated at 1200 cells/μl and 1100 cell/μl for male and female samples, respectively with greater than 80% cell viability. Single-cell capture, library preparation was done using a Chromium Next GEM Single Cell 3’ GEM, Library and Gel Bead Kit v3.1 on a 10X Chromium controller (10X Genomics). The volume of suspension to be loaded was decided as per the manufacturer recommendation to ensure the target capture of 6000 cells per well. Single cell suspension from male and female samples was loaded onto two separate wells of different chips giving a total of four libraries (2 male and 2 female samples). Each library was barcoded and sequenced separately on a single lane of HiSeq X to obtain a mean depth of at least 100,000 reads per cell in 2 x150bp configuration.

### Single cell RNA sequencing data analysis

#### Trimming the reads

To obtain informative portion of raw reads, they were hard trimmed to make sure read-1 and read-2 are 28 and 91 bps long as per requirements specified by 10X genomics using *trim_galore*.

#### Alignment, UMI counting and cell calling

Mouse reference genome mm10 (Mus_musculus.GRCm38.dna.primary_assembly.fa) and GTF file were downloaded from Ensembl (Genome build GRCm38.p6). The GTF file was filtered to retain only protein coding transcripts by removing readthrough and any non-coding transcripts using *cellranger mkgtf* using attribute=gene_biotype:protein_coding and readthrough_transcript. A custom reference was built with *cellranger mkref* using filtered GTF and mm10 genome. Alignment to custom reference, UMI counting and cell calling, removal of empty droplets and count matrix generation was done in a single step using *cellranger count* individually for each sample using default parameters.

#### Integrating samples and read depth normalization

After alignment, integration of 2 male and 2 female samples was done at raw data level ensuring average number of confidently mapped reads for each sample are equal using *cellranger aggr* pipeline. This pipeline subsampled reads in higher depth samples to ensure the sequencing depth is normalized across samples. During this step, a sample suffix (1 to 4) was added to each cell barcode to distinguish its source and to avoid barcode clashes in the integrated count matrix.

#### Ambient RNA correction, quality check and filtering

Before downstream analysis stringent filtering was implemented to retain high quality cells. Ambient RNA contamination was removed using soupX package with default parameters in auto mode. The adjusted count matrix from soupX consisting of 10615 cells was used as an input to Seurat package using *CreateSeuratObjec*t function. To remove potential doublets and low gene count cells, an additional filter was applied to select cells that express 200-7000 genes resulting in dropping 683 cells.

#### Normalization, scaling, dimensionality reduction, clustering, marker identification and additional filtration

The data was normalized using *NormalizeData* function using LogNormalize method that log-transforms the expression counts after multiplying with scaling factor 10000. After normalization, top 2000 variable genes in the dataset were identified using *FindVariableFeatures* function using mean.function=ExpMean, dispersion.function=LogVMR, x.low.cutoff =0.0125, x.high.cutoff=3, y.cutoff=0.5 as parameters. This variable gene set was used for all downstream analysis including dimensionality reduction and clustering. All vomeronasal receptors were amongst the variable features gene set. Data was scaled and centered using *ScaleData* using default model by regressing out percentage of mitochondrial genes and total number of RNA molecules per cell to remove their contribution in downstream analysis. The basis of scaling for each gene is by subtracting the mean from the value and dividing the difference by standard deviation. To cluster the cells, we initially performed principal component analysis (PCA) with 50 components using *RunPCA*. We used elbow plot and Jack Straw plot and determined the optimum number of dimensions required for the next steps of clustering as 37. Graph based clustering was performed using Seurat’s *FindNeighbours* function to identify the neighboring cells that share similar expression pattern in a network constructed in PCA space with 37 dimensions. Later the clusters were identified by using *FindClusters* function by varying the resolution parameter from 0.2 to 0.8. The resolution parameter 0.3 was chosen, as it shows minimum overlapping markers upon plotting the heatmap of gene markers for each cluster identified by *FindAllMarkers* function. To clean up the dataset further, cells (408) expressing Hbb-bs gene were considered as RBC contamination and a cluster (215 cells) enriched with mitochondrial genes indicative of dying cells were removed from the data. This resulted in a total of 9180 cells.

#### Dimensionality reduction for 2D representation and cell type assignment

UMAP was generated using *RunUMAP* function. Cluster identity was assigned based on known markers of each cell type. Two clusters expressing Gnai2 as major marker were merged as they had very similar expression profile. Solitary chemosensory cells and endothelial cells were manually assigned a cluster identity based on the expression of Trpm5/Rgs21 and Aqp1/Egfl7 and by selecting the cells using the *CellSelector* function of Seurat.

#### Comparison of male and female data

Based on suffix assigned to barcodes during integration of male and female samples, a metadata column was added to the Seurat object marking the sex of source tissue as male or female. Further, cells in each cluster were divided by appending the sex to the cluster identities. Differential expression analysis was performed between cells from male and female VNO for each cluster using *FindMarkers* function of Seurat.

#### Clustering and downstream analysis of neurons

For neuron specific analysis, neuronal cell types representing Gnao1 neurons, Gnai2 neurons, immature neurons, and progenitor cells were separated from main seurat object using *subset* function to create a new ‘neurons’ object. The data is scaled again, PCA was performed, number of principal components were determined, and clustering was performed as described above with resolution=0.4 to define neuronal subtypes. To compare the results of clustering with and without VRs, a new Seurat object– ‘neurons_no_VR’ was created by removing all vomeronasal receptors from the variable gene set. Dimensionality reduction, clustering was performed again using the same method mentioned above. Seurat’s query to reference mapping module was used to project neurons_no_VR object in the same UMAP space as neurons object so that UMAPs are comparable. The anchors were identified by using *FindTransferAnchors* function with neurons object as reference and neurons_no_VR as query. The reference UMAP model was computed using *MapQuery* function using the anchors.

#### Pseudotime analysis

Developmental trajectory of VSNs was inferred using SlingShot package. ‘neurons’ objects with 5 major clusters: progenitor cells, Gnao1+ immature neurons, Gnai2+ immature neurons, mature Gnao1 neurons and mature Gnai2 neurons. Progenitor cells/cluster n5 was chosen as starting cluster.

#### Differential gene expression of Gnao1, Gnai2 neurons

Differentially expressed genes between Gnao1 and Gnai2 neurons (mature or immature) were identified using *FindMarkers* function of Seurat on ‘neurons’ object with default parameters. The results are plotted as volcano plot using EnhancedVolcano package.

#### Gene ontology (GO) analysis

Gene ontology analysis of Gnao1 enriched genes was done with clusterProfiler package using *enricher* function on mouse gene sets related to mouse biological processes (GO:BP) ontology downloaded from Mouse Molecular Signatures Database. The enrichment analysis was restricted to the differentially expressed genes with log2 fold change greater than 1.

#### Co-expression analysis

Non-zero expression level for a particular gene may not indicate that the VR is expressed. Therefore, we identified the cut-off for V1R, V2R and H2-M*v*s based on the distribution of expression level across all cells. Normalized gene expression values of V1R, V2R and H2-M*v* genes were extracted from each cell of Gnai2 and Gnao1 clusters and the distribution is plotted across all cells. This resulted in a Bi-modal distribution and starting of the second peak was considered as cut-off for V2R/V1R/H2- M*v* genes (**Figure 4-figure supplement 1D, 1F**). The genes were considered co-expressed in a single cell if their normalized expression value is greater than the identified threshold value of 1.25 for V2R, H2-M*v* gene and 2.5 for V1R genes. Further, the number of cells crossing the threshold were calculated for each combination.

Raw single cell RNA seq data of VNO from p56 animals generated by (Hills *et al*., 2024) was downloaded from NCBI GEO with ID: GSE252365. The data was analyzed in the same method as described above and Gnao1 neurons from mature neuronal clusters identified based on the expression of Gnao1, Omp were used for co-expression analysis.

The graphical user interface for scvnoexplorer.com was made using ShinyCell tool (Ouyang *et al*., 2021). Reference details of software used for data analysis are mentioned in a tabular form below.

### RNA *in situ* hybridization (ISH)

**Probe design and synthesis**: Primers **(Supplementary table 11)** targeting unique regions of each gene were designed by adding T7/SP6 promoter sequence. PCR (GoTaq DNA polymerase) was performed using gene specific primers with VNO cDNA as template and the product was run on agarose gel to confirm specific amplification of each gene. When multiple bands were seen, the band with desired molecular weight was cut from the gel a. PCR product clean-up or gel purification was performed using NucleoSpin Gel and PCR Clean-up kit. Purified PCR product was verified by sanger sequencing and was used as template for in vitro transcription to obtain digoxigenin (Dig) and fluorescein (Flu) labeled riboprobes using following reaction composition: 1ug Template DNA, 1x Dig/Flu RNA labeling mix, 1U/uL RNAsin plus, 5mM DTT, 1x Transcription buffer in nuclease-free water. The reaction was purified using Qiagen/MN RNA cleanup kit and stored at -80℃ by adding formamide to 50% of volume.

H2-M*v* genes and Gnai2 were cloned to a plasmid vector with T7 or SP6 promoters. Gnao1 plasmid was purchased from Invitrogen (6309166). The plasmid was linearized, and in vitro transcription was performed as described earlier.

### Chromogenic ISH

Fresh VNO was embedded in OCT and 14µm thick sections were collected on a cryostat. Tissue sections were fixed with 4% PFA in Diethyl Pyrocarbonate (DEPC) treated phosphate buffer saline (PBS) and acetylation was performed using Propionic Anhydride, Triethanolamine, NaCl. Permeabilization was done using 0.1M HCl and sections were pre-hybridized using hybridization buffer (50% Formamide, 5X SSC, 5X Denhardt’s solution, 0.1mg/mL Salmon sperm DNA, 0.25mg/mL yeast t-RNA) for 2 hours. Probes were diluted (1:100) in the hybridization buffer and added to the slides. Hybridization was performed for 12-16 hours at 67°C. To remove the unbound probe, sections were washed thrice with 0.2X SSC for 30 minutes each. Slides were equilibrated in a buffer consisting of 0.1M Tris-HCl. pH 7.5, 150mM NaCl for 5 minutes and blocked with 10% FBS in the same buffer. Hybridized Digoxigenin (DIG) or Fluorescein (Flu) containing RNA probes were detected by alkaline phosphatase conjugated Anti-DIG or Anti-Flu Fab2 fragments (1:7500 dilution). Unbound antibody was washed, and development of alkaline phosphatase was done using BCIP/NBT substrate diluted as per manufacturer’s protocol in 0.1M Tris-HCl pH-9, 0.1M NaCl, 50mM MgCl2. The signal was developed for 12 - 72 hours based on the intensity. After development, slides were washed with PBS and mounted using 10% glycerol in 0.1M Tris-HCl (pH 7.5).

### Two-color fluorescence ISH

As described above, the same protocol for chromogenic ISH was followed until 0.2X SSC washes, after which the hybridized probes were detected using peroxidase conjugated anti-FLU or anti-DIG Fab(2) fragments and the Tyramide signal amplification (TSA) system from Akoya Biosciences. Slides were incubated in 3% H2O2 in PBS for 1 hour and washed thrice for 10 minutes each and blocked with 0.5% blocking buffer for 30 min. Dig and Flu were sequentially developed using TSA-FITC or TSA-Cy3.

### Immunohistochemistry

VNOs were dissected from animals of age 8-12 weeks, fixed with 4% Paraformaldehyde (PFA) in PBS and cryopreserved with 30% sucrose. Tissue was embedded in OCT- freezing medium and cryosections of 14µm thickness were collected on glass slides. Sections were post fixed again with 4% PFA in PBS, blocked and permeabilized by incubating for 2 hours with a blocking buffer (3% Bovine Serum Albumin, 0.1% TritonX- 100 in PBS with 0.02% Sodium Azide). After blocking, the sections were incubated with primary antibodies diluted in the blocking buffer for 2 hours followed by washing off excess antibody and secondary antibody incubation for 2 more hours. All washes were done using 0.1% TritonX-100 in PBS.

### Light microscopy image acquisition

Chromogenic images were acquired on Olympus BX43 upright microscope using bright field Koehler illumination and 10X objective (PlaN, 0.24 NA) equipped with Olympus DP25 color CCD camera. Two or three color fluorescent RNA-ISH images and fluorescent Immunohistochemistry images were acquired sequentially using Leica Stellaris upright confocal microscope using 10X (HC PL APO 0.40 NA) or 63X (HC PL APO 1.40-0.60 NA Oil) objective keeping 1 Airy unit as pinhole size. Acquisition parameters were adjusted and calibrated using single channel controls to ensure that there was no spectral cross-talk between channels in multi-color imaging experiments.

### Quantification of signal intensities of ER antibodies

VNO sections were labelled with a single ER antibody, along with goat anti-OMP and rabbit anti-Gnao1 or mouse anti-KDEL as markers of neurons and Gnao1 zone respectively. At least 20 sections from 3 replicates were imaged via confocal microscopy and used for each ER antibody quantification. 50-pixel wide rectangular regions of interest (ROI) were drawn along apical to basal axis and the signal intensity along the ROI for each channel were extracted from the images using Fiji software. To make the signal intensities along apical-basal axis across different images and antibodies comparable, the signal intensity and length of ROI were normalized by min-max normalization

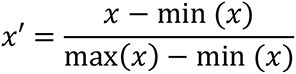

where, 𝑥 is the measured or actual value, min(𝑥) and max(𝑥) are minimum and maximum values of 𝑥, respectively. The trendline of normalized signal intensity along the ab axis was generated by fitting a smoothened curve using *geom_smooth* function of ggplots package based on generalized additive model for each antibody.

### Electron Microscopy

VNOs were dissected from animals that were trans-cardially perfused with PBS followed by 4% PFA in PBS buffer. Dissected VNO was embedded in 2% agarose and coronal sections of 400um were obtained on a vibratome in 0.1M sodium cacodylate buffer (pH 7.4). Sections were fixed with Karnovsky’s fixative (3% PFA + 2% glutaraldehyde in 0.1 M sodium cacodylate) for up to 1 week. Subsequent processing steps were similar to that described before (Terasaki, Brunson and Sardi, 2020). Briefly, vibratome sections were rinsed with 0.1 M sodium cacodylate then immersed in 1% osmium, 0.8 % potassium ferricyanide in the cacodylate buffer for 1 hour. This was followed by incubation in 1% aqueous uranyl acetate 1 hr, then 30 min in lead aspartate (Walton, 1979), dehydration in graded ethanol, infiltration with epon resin then embedding and polymerization at 60 °C for 2 days. Serial 70 nm thick sections were collected using a Powertome automated tape collector (RMC Boeckeler, Tucson, AZ). The tape was mounted on a silicon wafer, carbon coated and imaged with a field emission electron microscope (Thermo Fisher / FEI Verios, Hillsboro, OR). Backscatter electrons were imaged from a 5 keV / 0.8 nanoAmp beam of electrons.

## Data availability

All raw data related to single cell RNA sequencing was deposited to GEO and can be publicly accessed using accession ID GSE253252.

## Acknowledgements

We acknowledge Jyoti Rohilla, Nandana Nanda for help in preparation of DNA templates for riboprobe generation, Tulasi Nagabandi for helping with scRNA library preparation, Tamal Das for sharing Atlastin1, Sec61β antibodies and Erik Snapp for insightful discussions. We acknowledge support from the Department of Atomic Energy, Government of India, under Project No. RTI 4007.

---

**Figure 1-figure supplement 1.**
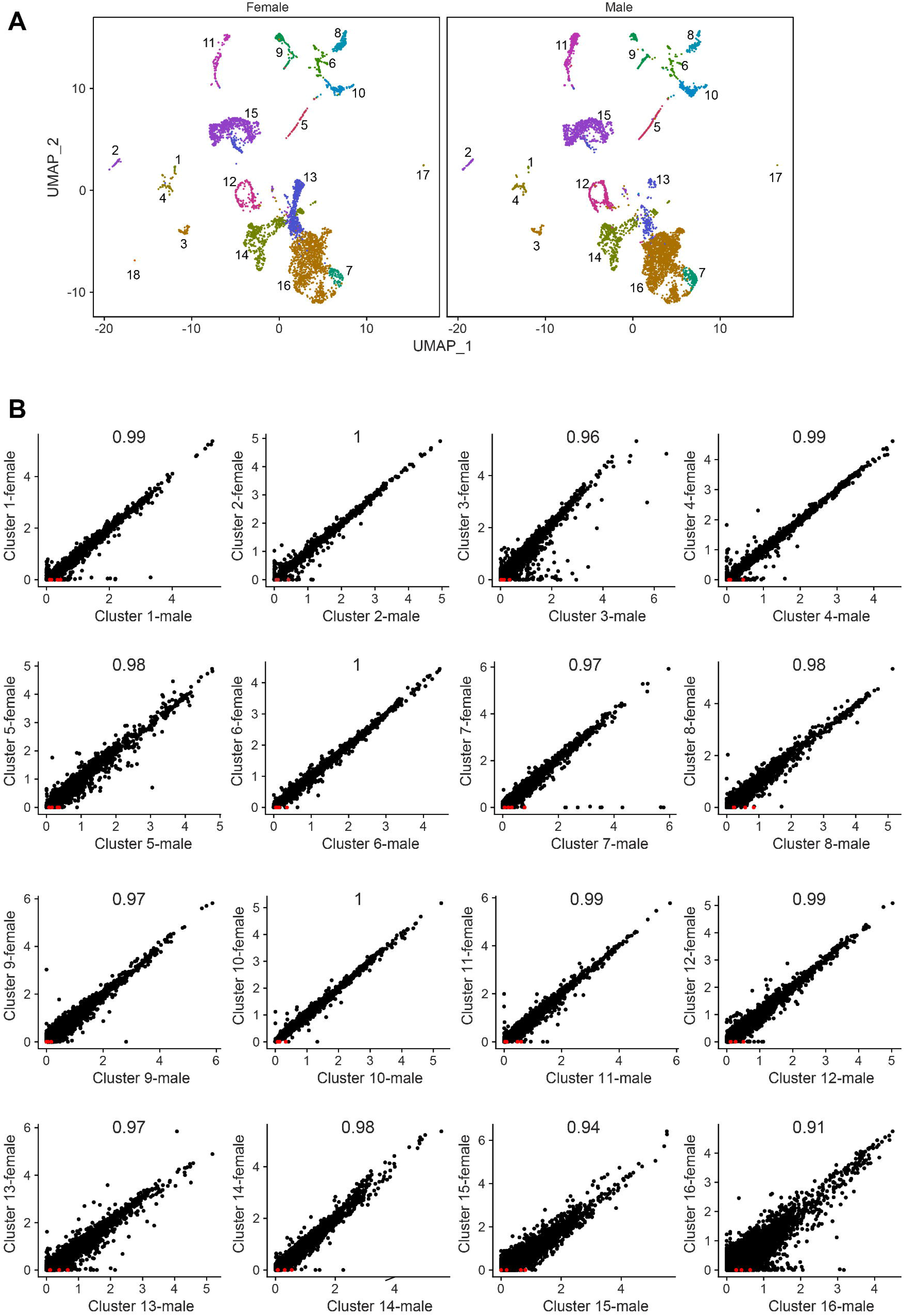
Comparison of cell type composition and gene expression from male and female vomeronasal neuroepithelium. **A)** Uniform Manifold Approximation Projection (UMAP) of cells from male and female vomeronasal neuroepithelium. with the cluster numbers corresponding to Figure 1A, 1**B.** Solitiary chemosensory neuron (Cluster 18) were seen only in female data. **B)** Scatter plots comparing average expression of genes accross each cluster from male and female with Pearson correlation co-efficient at the top of the plot. Each point in the plot represents a gene. Known sexually dimorphic genes: Eif2s3y. Ddx3y, Uty. Kdm5d are marked in red. Scatter plot of cluster 17 between male and female is not shown due to low cell count. The results of differential expression between each cluster of male and female are in Supplementary table 1.

**Figure 3-figure supplement 1.**
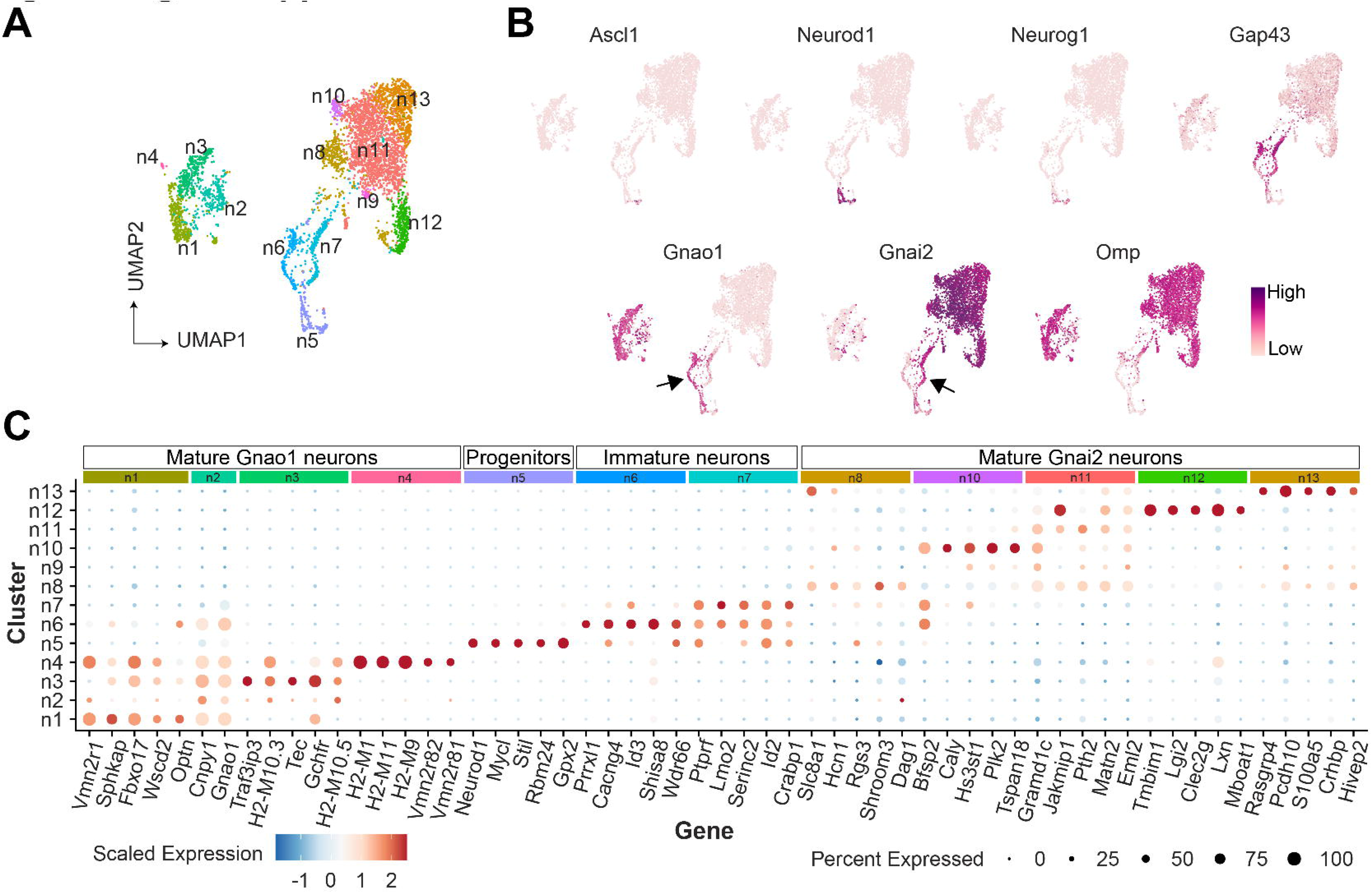
A) UMAP projection of neurons with 13 clusters (n1-n13). B) Feature plot showing expression of neuronal markers associated with various stages of differentiation: Globose basal cells (n5; Ascl1), progenitors cells (n5; Neurod1, Neurog1), immature neurons (n6, n7; Gap43+), mature Gnao1 neurons (n1-n4; Gap43-, Omp+, Gnao1+) and mature Gnai2 neurons (n8-n13; Gap43-, Gnai2+, Omp+). Arrows highlight the expression of Gnao1 and Gnai2 in Gap43+ immature neurons (n6, n7). C) Dotplot showing enriched genes in each cluster compared to all other clusters obtained by differential expression analysis. Size of the dot represents the percentage of cells expressing the gene in that cluster and colour indicates the scaled expression value. Top five gene markers based on log2fold change from each cluster were chosen by filtering the genes whose adjusted p-value is less than 0.005, expression in atleast 50% of cells and less than 50% of cells of all other clusters. No markers were found for cluster n9 with this filtering criteria. Complete list of top 30 markers for neuronal clusters is in Supplementary table 5.

**Figure 3-figure supplement 2.**
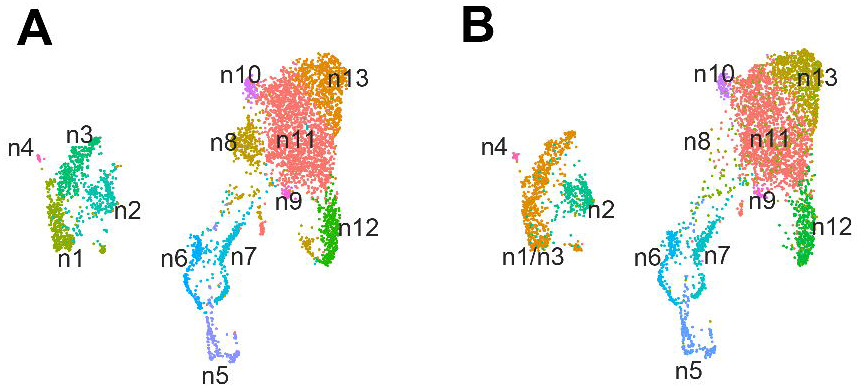
**Effect of VR genes on neuronal clustering.** Clustering of neurons based on top 2000 variable geneset in the dataset leading to 13 clusters (same as Figure 3**-figure supplement 1A).** Reclustering performed after excluding genes coding vomeronasal receptors from the variable geneset without changing any other parameters. The bifurcation of Gnao1, Gnai2 neurons at mature and immature stages remains unchanged. The only changes are merger of mature Gnao1 sub-clusters n1/n3 and change in mature Gnai2 n8 composition.

**Figure 4-figure supplement 1.**
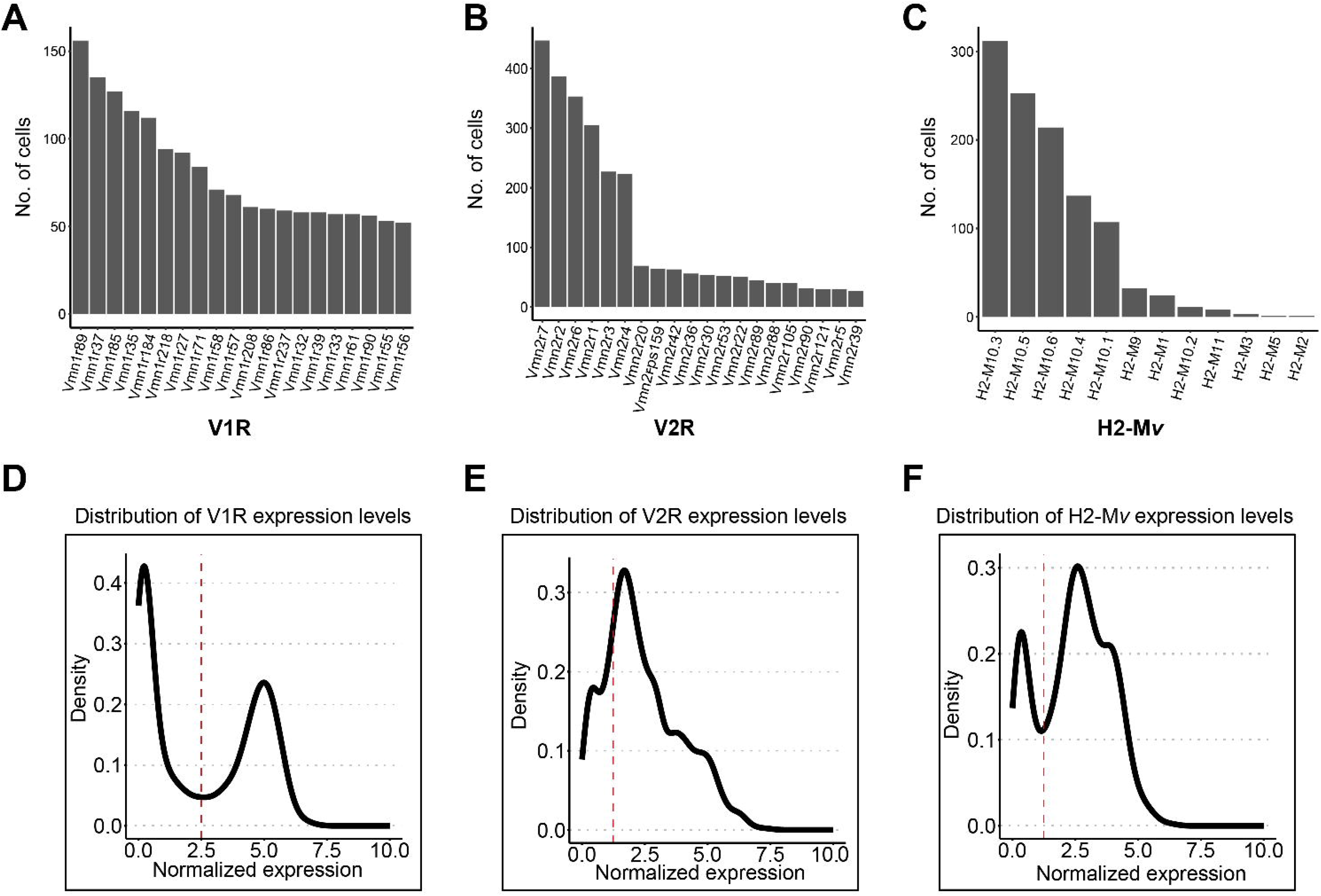
Bar plot showing the number of cells expressing top 20 V1Rs **(A),** V2Rs **(B)** and H2-Mvs **(C).** Normalized gene expression values were extracted for all V1Rs **(D)** from Gnai2 neurons, V2Rs (E) and H2-Mvs **(F)** from Gnao1 neurons and density was plotted to identiy the distribution of gene expression. The red dashed line represents a normalized gene expression value that was used as threshold to call **the expression of respective genes.**

**Figure 4-figure supplement 2.**
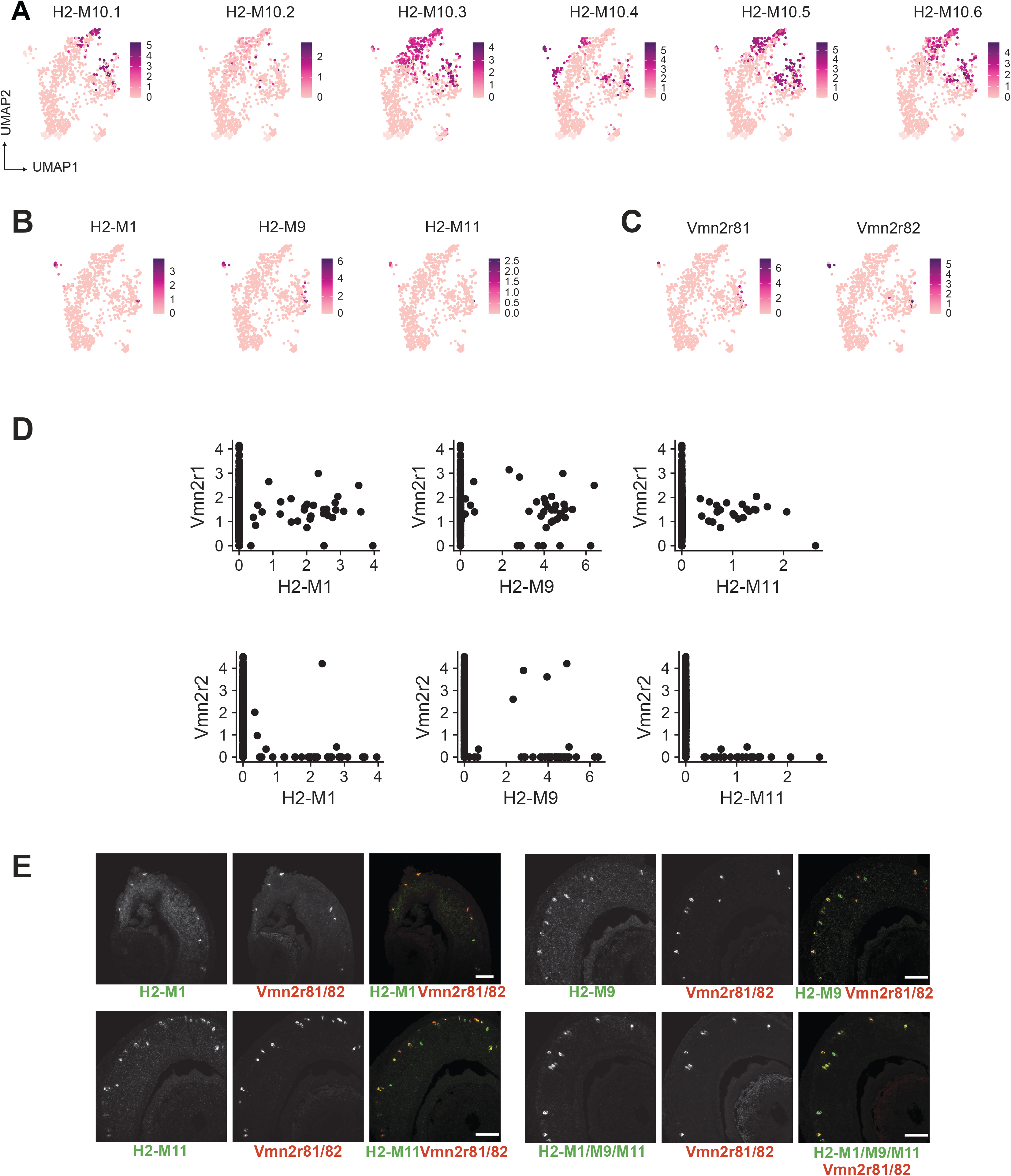
Characteristics of H2-M*v* expression **A -B)** Feature plot showing the expression H2-M10 family genes **(A)** and limited expression of phylogentically divergent H2-M*v* members H2-M1, H2-M9 and H2-M11 **(B)** in Gnao1 neurons. **C)** Feature plot showing the expression of Vmn2r81 and Vmn2r82 in few cells of cluster 2 and 4 of Gnao1 neurons that express H2-M1, M9 and M11. **D)** Scatter plot showing the normalized expression level per cell of rarely expressed H2-M*v* genes (H2- M1, M9 and M11) on x-axis and Vmn2r1 or Vmn2r2 on y-axis indicating that H2-M1, M9 and M11 co-express with Vmn2r1 unlike other H2-M*v* genes**. E)** Two-color RNA in situ hybridization of H2-M1,H2-M9 and H2-M11 with Vmn2r81/82 confirming the co-expression. The ISH probe was common for 9U81 DQG 82; +2-01/09/011 LQGLFDWHV SRROLQJ RI LQGLYLGXDO SUREHV. 6FDOH EDU: 100 ȝP.

**Figure 4-figure supplement 3.**
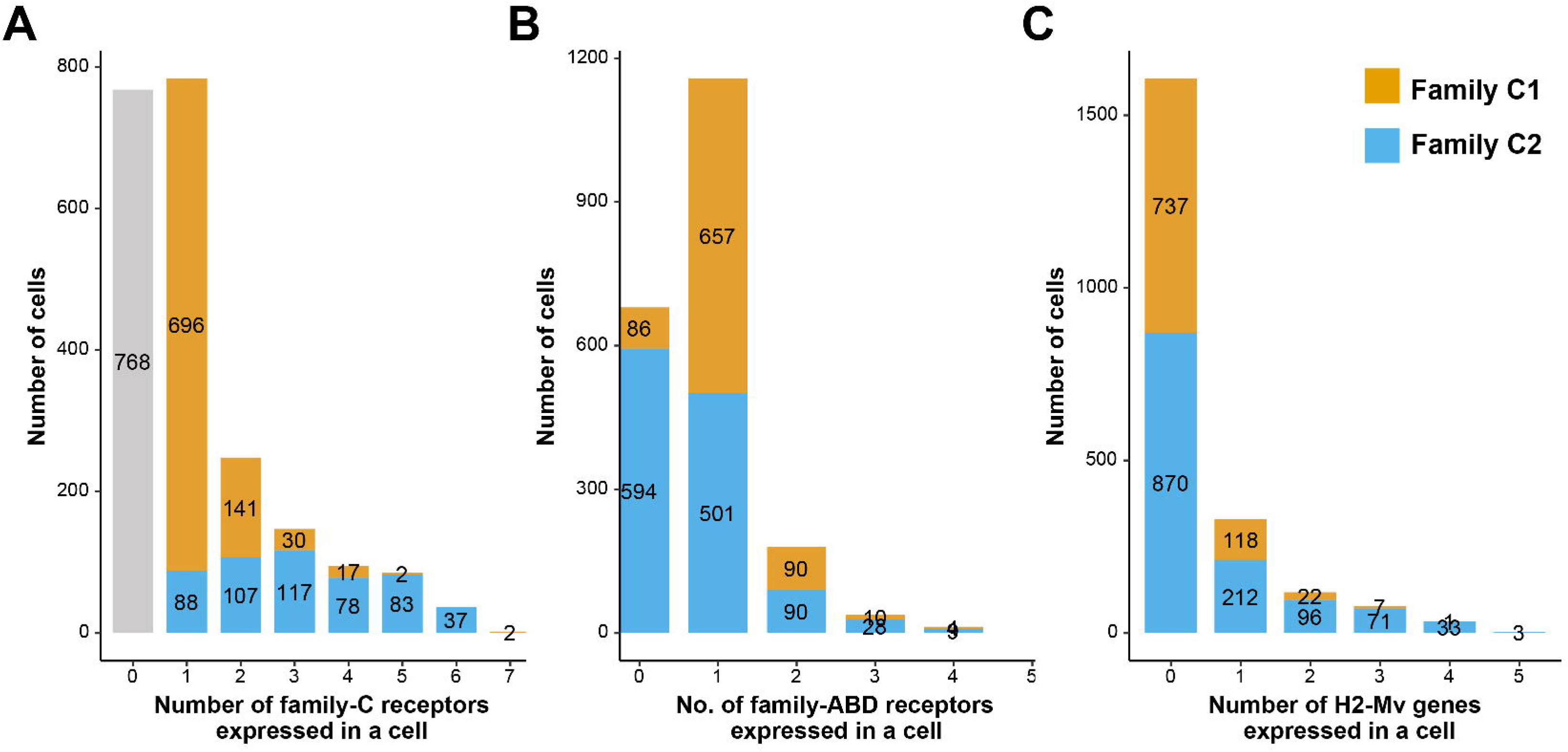
Coexpression characteristics of V2R and H2-Mv genes using data from Hills et al., 2024 **A-C)** Bar plot showing number of cells expressing: 0-7 family-C V2Rs per cell **(A),** 0-5 family-ABO V2Rs per cell **(8),** 0-6 H2-Mv genes per cell **(C)** with composition of cells associated with family C1 (orange) or C2 (blue) V2R color coded on the bar. The trend of co-expression is similar to Figure 7F, 7G and 7H indicating that co-expression counts are similar across datasets.

**Figure 4-figure supplement 4.**
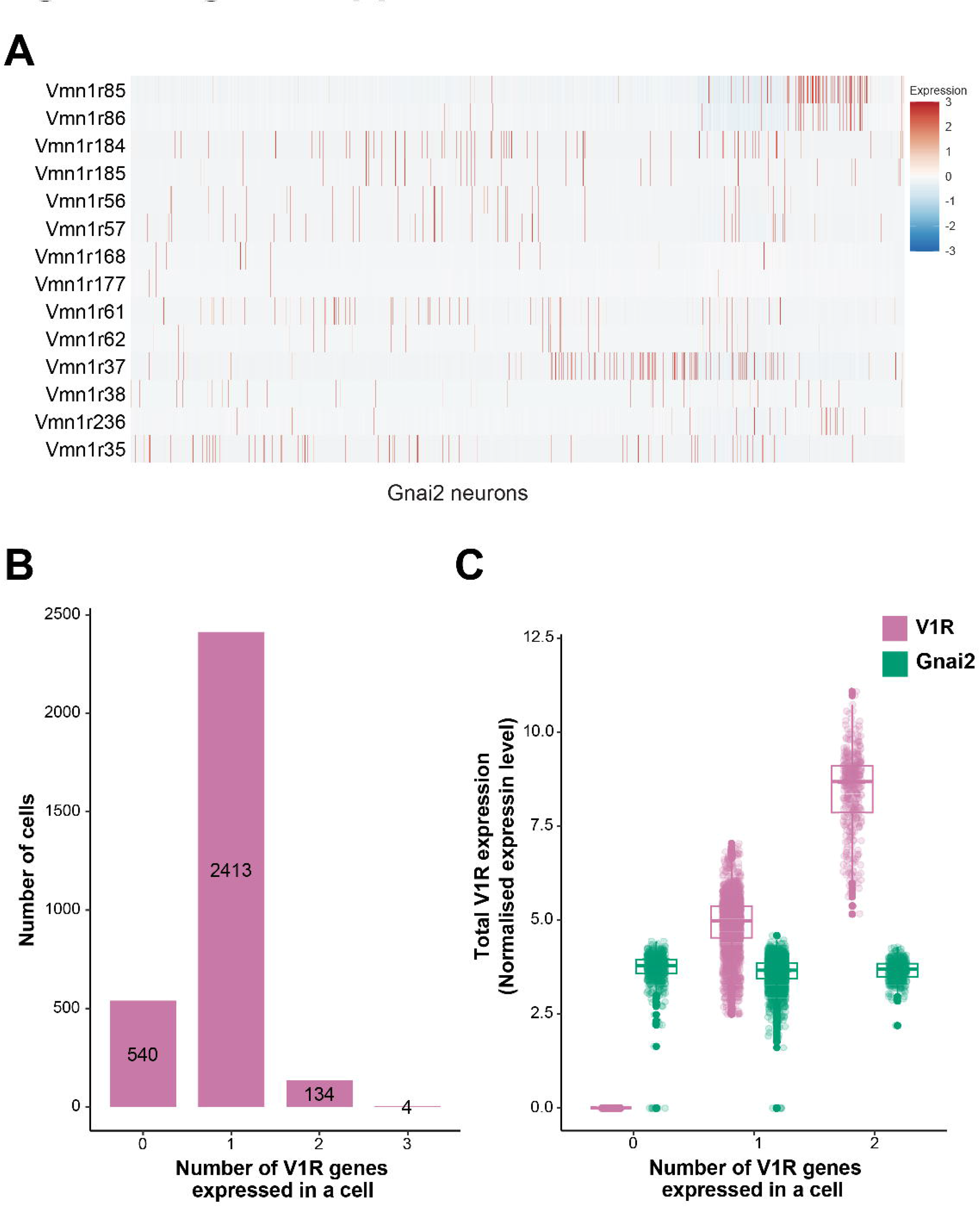
Co-expression of V1Rs in Gnai2 neurons A) Heatmap showing the expression of selected V1Rs in Gnai2 neurons. Each column represents a cell and the scaled expression value of each VR is color coded in each row with red and blue indicating high and low expression respectively. A continues red line in two rows of a single column indicates the expression of two receptors in a single cell. B) Bar plot showing number of cells expressing 0-3 V1Rs per cell indicating the V1R co-expression is limited to small subset of cells. **C)** Box plot comparing total V1R and Gnai2 (green) expression from cells shown in **B.** Multiple combinations of V1Rs co-expressed in a single cell and their cell frequency are listed in **Supplementary table 8.**

**Figure 5-figure supplement 1.**
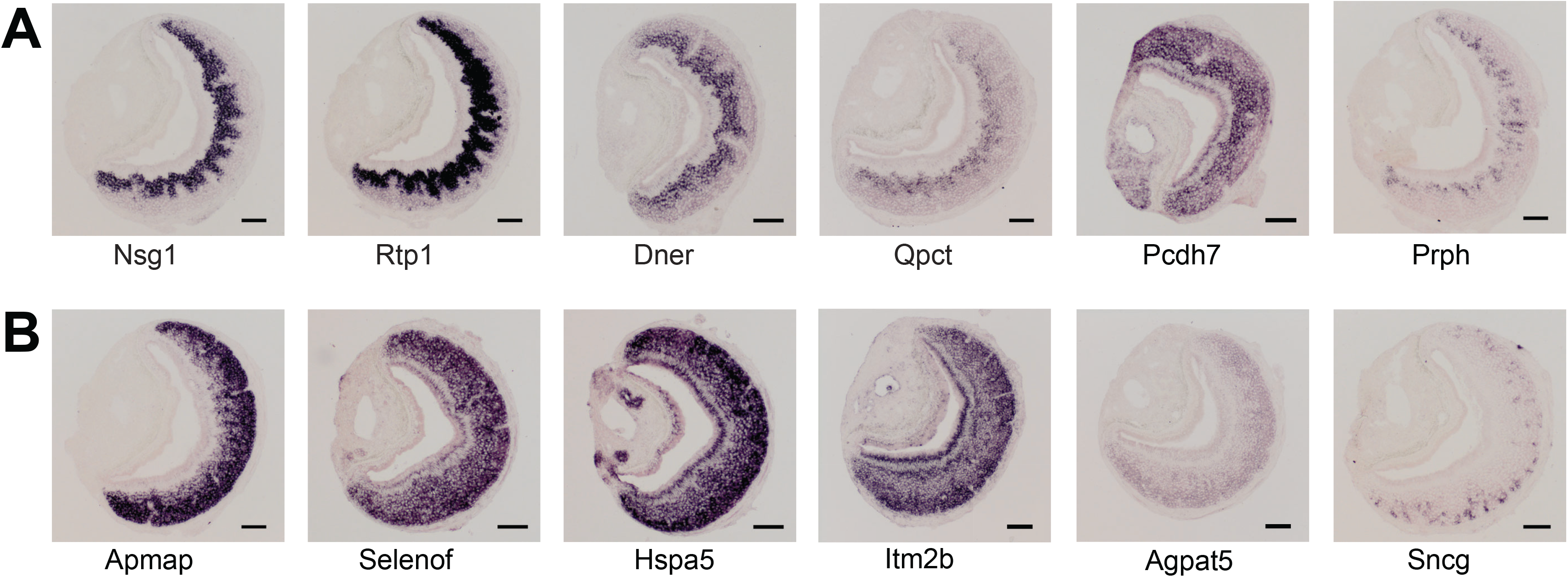
**Expression pattern of enriched genes in Gnai2 or Gnao1 neurons**. Chromogenic RNA-ISH showing expression of Gnai2 enriched genes: Nsg1, Rtp1, Dner, Qpct, Pcdh7, Prph (**A**) and Gnao1 enriched genes: Apmap, Selenof, Hspa5 (Bip), Itm2b, Agpat5, Sncg **(B)**. Sncg and Prph are expressed in a scattered pattern amongst few neurons. Scale bar: 100 μm

**Figure 7-figure supplement 1:**
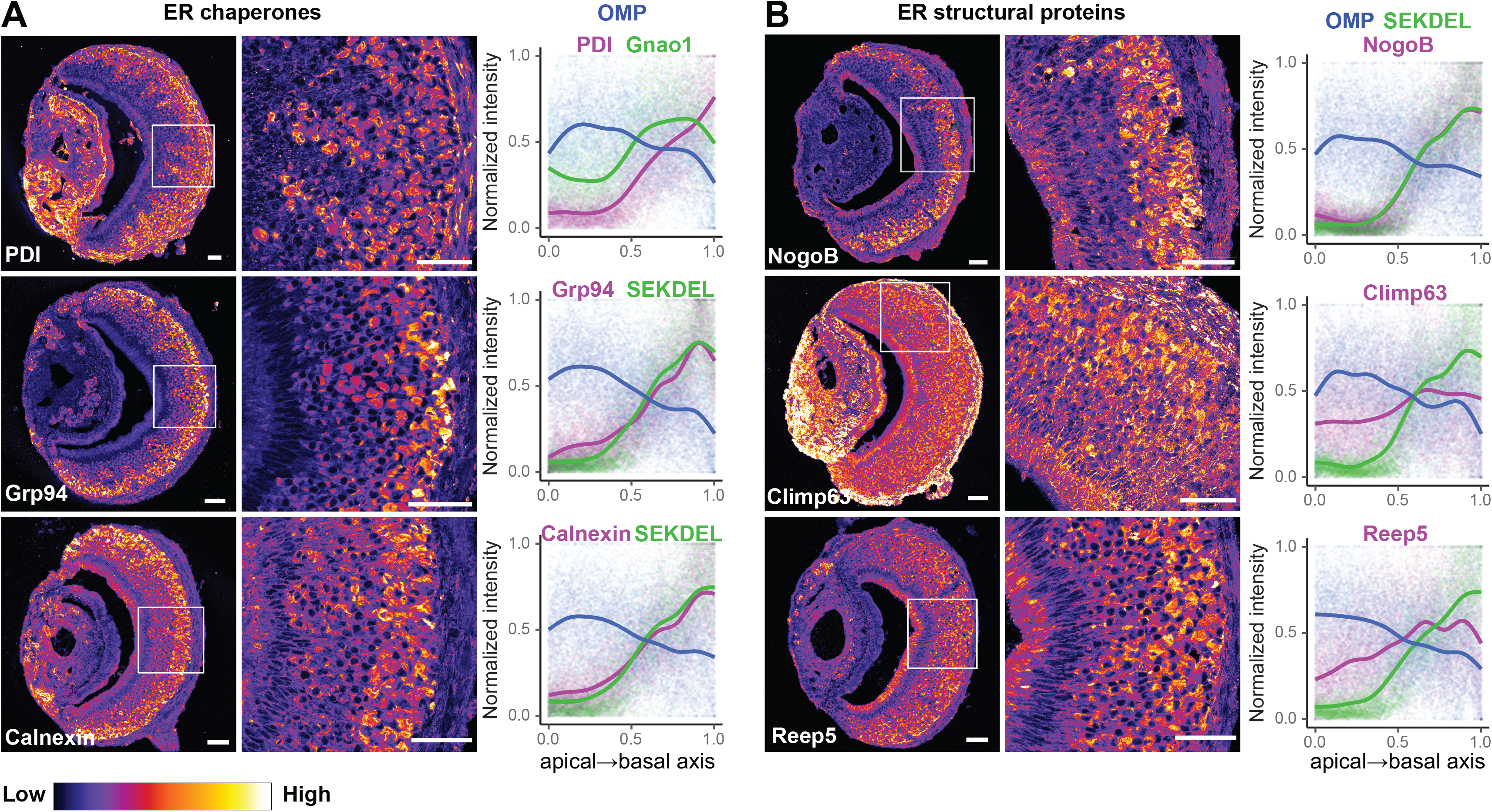
Pseudocolored immunofluorescence images of VNO cornonal sections labeled with antibodies against ER chaperone proteins: PDI / Grp94 / Calnexin (A), ER structural proteins: NogoB / Climp64 / Reep5 (B). Higher intensity in Gnaol neurons compared to Gnai2 neurons is seen from the images and quantified along the apical-basal axis by co-labelling sections with anti-OMP to mark all neurons and anti-Gnao1 (or anti-SEKDEL -see Figure 7A-E epending on antibody species), to mark Gnaol neurons. Fluorescence intensity along the apical-basal axis is quantified from atleast 20 sections from 3 nimal replicates. The normalized fluorescence intensity of ER proteins increases along apical-basal axis, similar to Gnaol or SEKDEL, while the level of MP either remains same or slightly decreases. Scale bar: 50 pm

**Figure 7-figure supplement 2.**
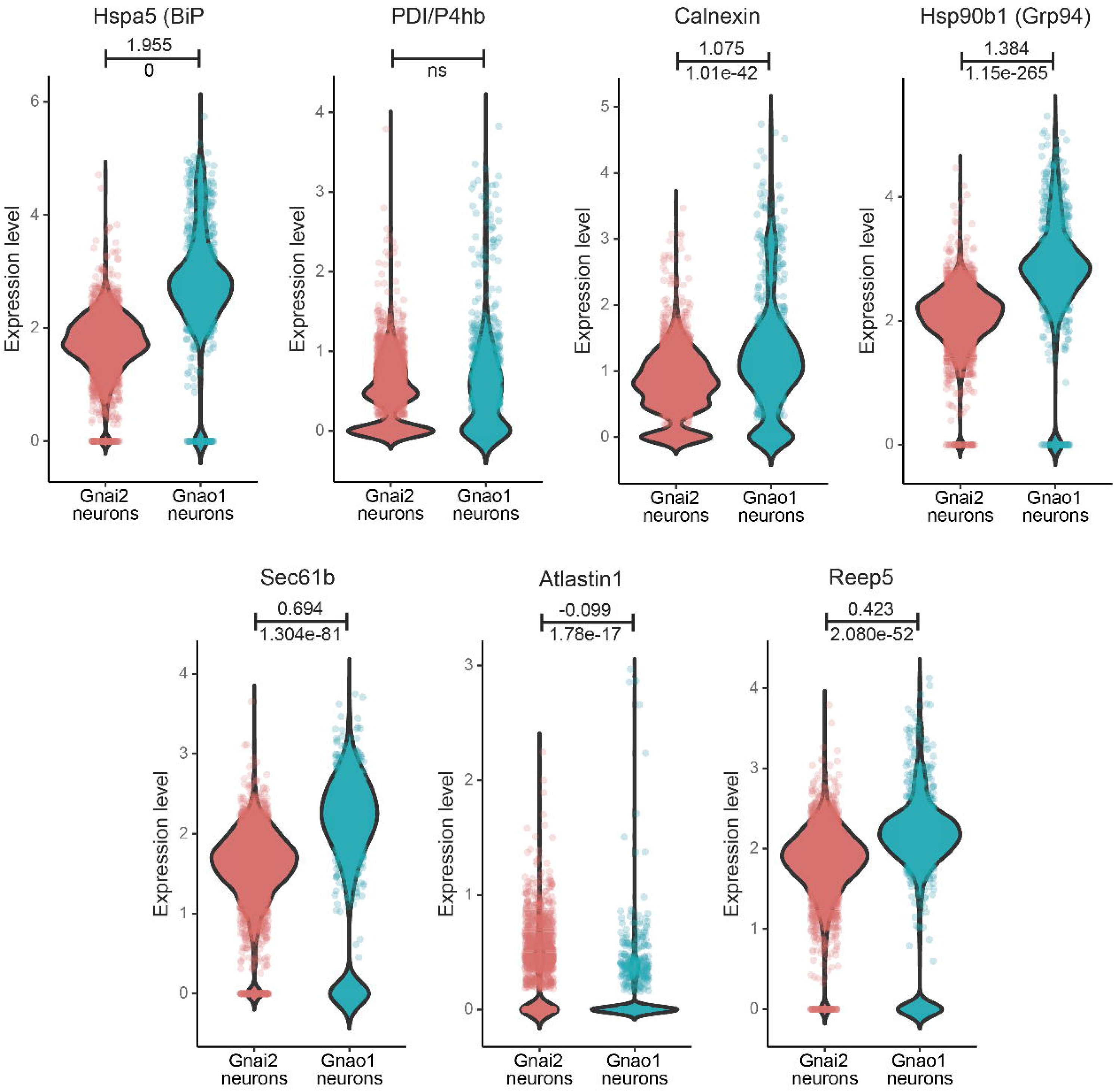
Comparison of ER gene expression between Gnai2, Gnao1 neurons. Violin plots showing the gene expression levels in mature Gnao1 or Gnai2 neurons whose protein levels are shown via immunofluorescence in Figure 7 and Figure 7-figure supplement 1. Log, fold change value for Gnao1 vs Gnai2 calculated from pseudobulk differential gene expression analysis is mentioned on top of the line and Bonferroni-adjusted p-value is mentioned below the line (ns - not significant). RNA levels of Hspa5. Calnexin. Hsp90b1. Sec61b. Reep5 are significantly upregulated in Gnao1 neurons while POI and Atlastin1 do not differ significantly between Gnao1 and Gnai2 neurons.

**Figure 8-figure supplement 1.**
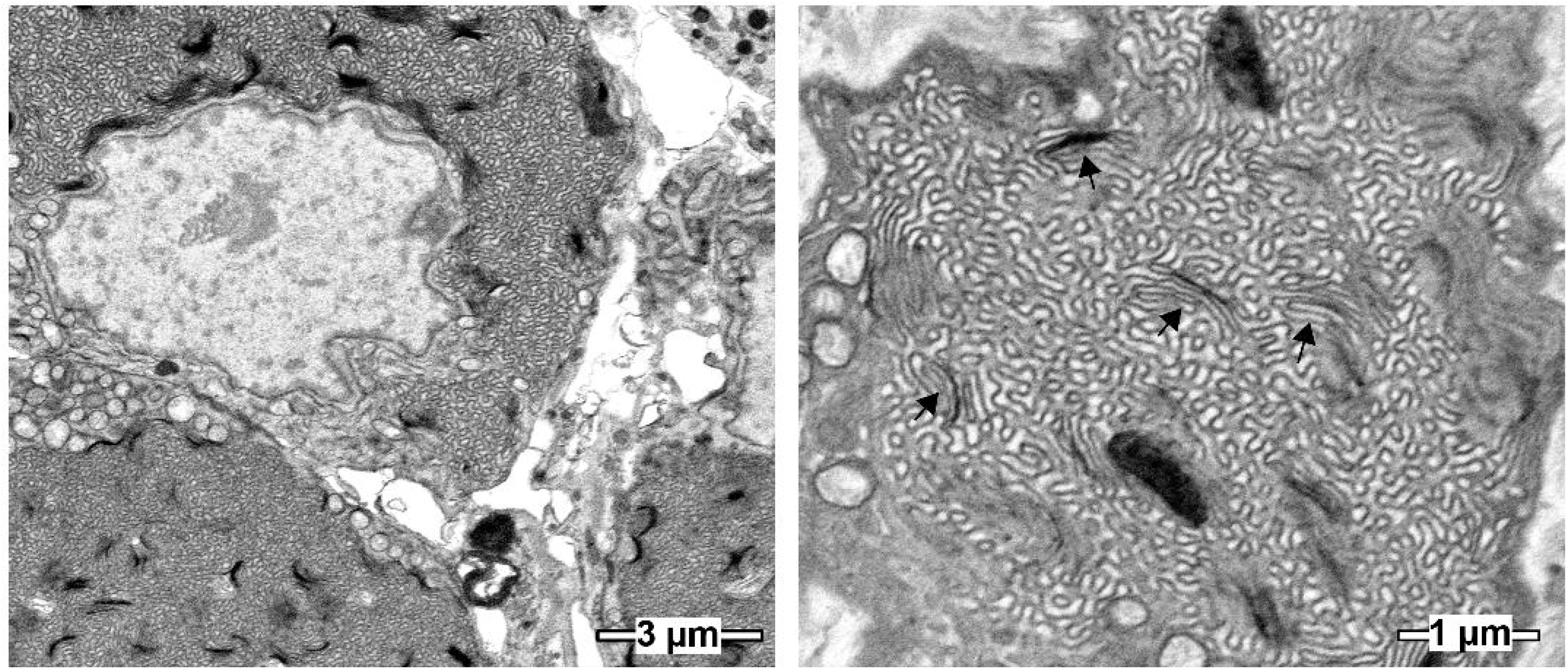
Scanning electron micrographs showing cell bodies of basal/Gnao1 neurons with sinusoidal ER membranes interspersed with stacked membranes (arrows).

